# The chromokinesin Kid (KIF22) forms a homodimer, moves processively along microtubules and transports double-stranded DNA

**DOI:** 10.1101/2024.03.13.584902

**Authors:** Shinsuke Niwa, Natsuki Furusaki, Tomoki Kita, Yuki Suzuki, Kyoko Chiba

## Abstract

During prometaphase in mitosis, chromosomes are pushed toward the spindle equator. The chromokinesin Kid, also known as KIF22, moves chromosomes along spindle microtubules during prometaphase. Kid has long been considered a monomeric and non-processive motor, different from typical kinesins. In this study, we demonstrate that the full-length Kid forms a homodimer and moves processively along microtubules. A conserved coiled-coil domain within the stalk region of Kid is sufficient for homodimer formation and is required for the processivity of Kid. Furthermore, the neck linker and coiled-coil domains of Kid could add processive activity to the motor domain of KIF1A, suggesting that Kid contains a functional neck linker and dimerization capability, a prerequisite for the processivity of kinesin motor domains. The full-length Kid, containing a helix-hairpin-helix domain, can transport double-stranded DNA along microtubules in vitro. Alphafold3 prediction suggests that the dimerization of Kid stabilizes the association with DNA. These findings collectively suggest the reclassification of Kid as a processive and dimeric motor that transports DNA along microtubules.

## Introduction

Microtubules are reorganized to form a bipolar spindle as cells enter mitosis. In the prometaphase of mitosis, chromosomes are transported along the microtubules toward the spindle equator (Rieder et al., 1986). This process, known as chromosome congression, requires kinesin-4, kinesin-10 and kinesin-12 class of motor proteins (Bieling et al., 2010; Iemura and Tanaka, 2015; Wordeman, 2010). The mechanical forces that move chromosomes toward the spindle equator is called polar ejection forces (Brouhard and Hunt, 2005; Levesque and Compton, 2001; Rieder et al., 1986). The Kinesin-like DNA-binding protein (Kid), belonging to the kinesin-10 family and also known as KIF22, serves as a main molecular motor that transports chromosomes and generates polar ejection forces (Brouhard and Hunt, 2005; Levesque and Compton, 2001; Ohsugi et al., 2003; Thompson et al., 2022; Tokai et al., 1996; Wandke et al., 2012). Structurally, Kid contains a kinesin motor domain (Yajima et al., 2003), a coiled-coil domain (Shiroguchi et al., 2003), and a DNA binding domain (Tokai et al., 1996) (Figure 1A). In mitosis, the DNA binding domain of Kid binds along chromosome arms (Antonio et al., 2000; Funabiki and Murray, 2000; Levesque and Compton, 2001). Using the motor domain, Kid transports chromosomes (Bieling et al., 2010; Brouhard and Hunt, 2005).

**Figure 1.**
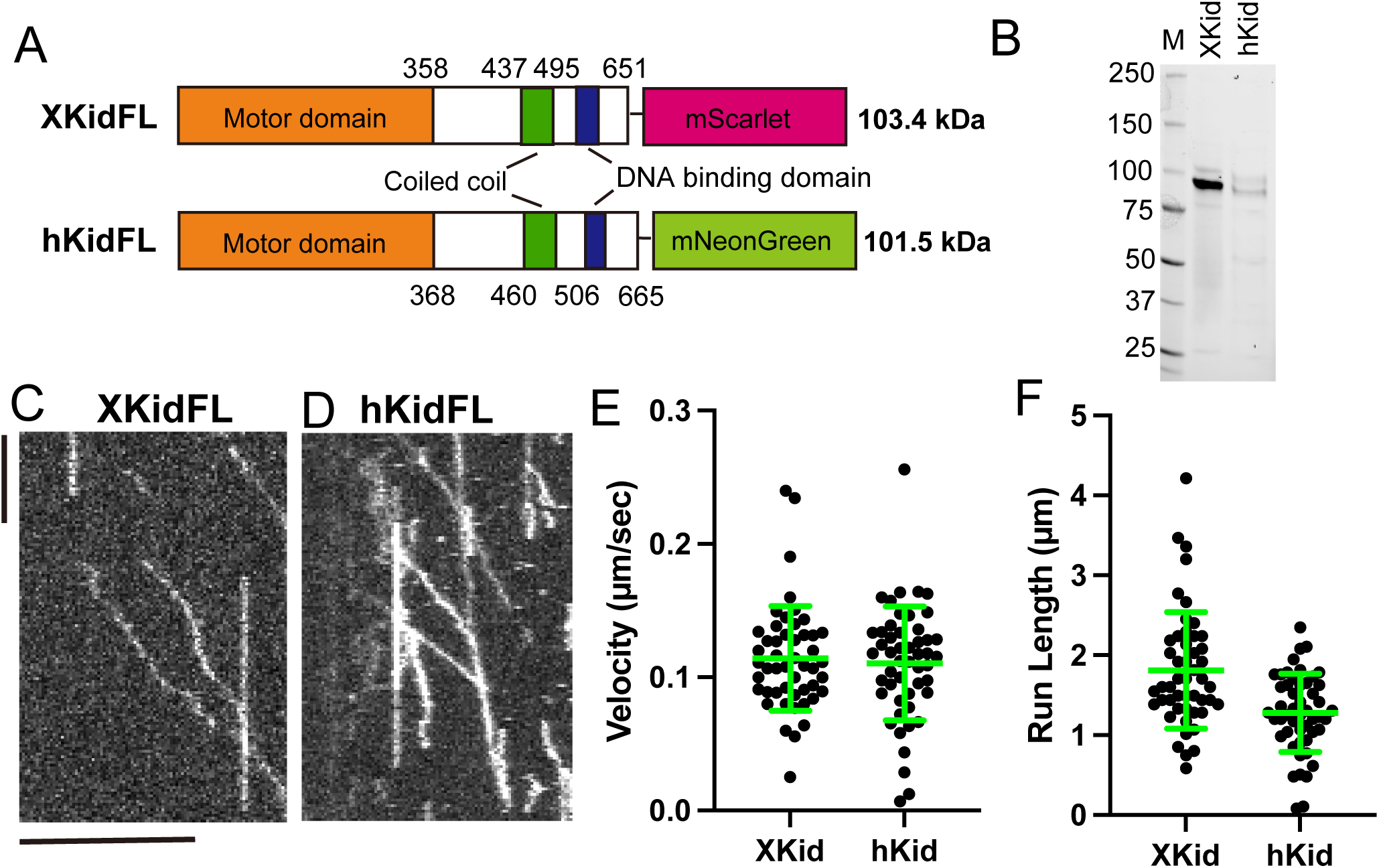
Kid is a processive motor. (A) Schematic illustration of the domain organization in Xenopus Kid tagged with a fluorescent protein mScarlet (XKidFL) and human Kid tagged with mNeonGreen (hKidFL). The calculated molecular weights of the fusion proteins are indicated on the right. (B) Representative SDS-PAGE analysis of purified XKidFL and hKidFL fusion proteins. The proteins are visualized using a Stain-Free gel. The molecular weight standards are indicated on the left side of the SDS-PAGE images. (C and D) Representative kymographs showing the motility of XKidFL at 20 pM (C) and hKidFL at 20 pM (D) both in the presence of 2 mM ATP. Scale bars: horizontal, 10 µm; vertical, 60 seconds. (E) Dot plots showing the velocity of XKidFL and hKidFL. Each dot shows a single datum point. Green bars represent mean ± S.D.. n = 51 and 52, respectively. (F) Dot plots showing the run length of XKidFL and hKidFL. Each dot shows a single datum point. Green bars represent mean ± S.D.. n = 51 and 52, respectively.

A single kinesin molecule can move for hundreds of steps along a microtubule without dissociating (Hackney, 1995; Hancock and Howard, 1998). This property, called processivity, requires dimerization of kinesins (Hancock and Howard, 1998). However, Kid has long been regarded as a monomeric and non-processive motor, which can move along microtubules in only a single step (Shiroguchi et al., 2003; Yajima et al., 2003). Consistent with these findings, Kid is unique within kinesin superfamily due to lack of the conventional neck coiled-coil domain (Shiroguchi et al., 2003), an element that determines the length of the neck linker essential for the coordinated movement of the two motor domains (Case et al., 2000; Isojima et al., 2010; Yildiz et al., 2008). On the other hand, a previous study showed that the coiled-coil domain of Kid interacts with full-length Kid in GST pull-down assays, suggesting this domain may mediate dimerization of Kid (Pike et al., 2018). In addition, full-length human Kid was reported to move processively along microtubules (Stumpff et al., 2012) and, Kid was proposed to be a chemically processive motor (Walker et al., 2019). However, these studies did not show the oligomeric state of Kid, nor did they provide detailed information on its motility characteristics and parameters. A Drosophila kinesin-10 motor NOD, an orthologue of Kid, is also characterized as a monomeric and non-processive motor (Matthies et al., 2001). Similar to Kid, NOD has a DNA binding domain at the tail domain (Afshar et al., 1995). NOD shows a processive movement only when the protein is forcedly dimerized by the addition of an artificial coiled-coil domain (Ye et al., 2018). Nevertheless, Kid acts as an active motor in microtubule gliding assays and cargo transport assays, both of which detect movement generated by multiple Kid motors (Bieling et al., 2010; Li et al., 2016; Shiroguchi et al., 2003; Takagi et al., 2013). Collectively, it has been widely assumed that Kid, unlike other kinesins, transports chromosomes by their cooperative action of many non-processive monomers (Brouhard and Hunt, 2005; Iemura and Tanaka, 2015; Takagi et al., 2013; Thompson et al., 2022).

Recently, we and others have analyzed properties of full-length kinesins (Chiba et al., 2022; Chiba et al., 2019; Fan, 2022; Wang et al., 2022). Notably, these studied have found that monomeric kinesins, including KIF1A, UNC-104 and KIF13B are converted to a dimer when activated (Chiba et al., 2023; Fan, 2022; Kita et al., 2024; Tomishige et al., 2002). For instance, in the autoinhibited and inactive state, KIF1A and UNC-104 are monomeric (Kita et al., 2024; Tomishige et al., 2002). The monomeric motor domain of UNC-104 and KIF1A has a plus-end directed motor activity but exhibits one-dimensional diffusion, meaning that the efficiency is low (Okada et al., 2003; Tomishige et al., 2002). Upon the release of autoinhibition, KIF1A and UNC-104 form dimers, which exhibit efficient directional movement on microtubules (Kita et al., 2024). These studies prompted us to reanalyze the oligomeric state and the motile properties of Kid, mainly in the full length. In this study, we show that the full-length Kid protein moves processively along microtubules. Kid proteins can form dimers but Kid proteins are dissociated to monomers at low concentrations. In the reconstitution assays in vitro, full-length Kid transport double-stranded DNA along microtubules.

## Results

### Kid exhibits processive movement along microtubules

To study biochemical and biophysical properties of full-length Kid, we firstly purified the full-length human Kid (hKid) and Xenopus Kid (XKid) using the baculovirus system and Sf9 cells because a previous study has succeeded in purifying functional XKid from Sf9 cells (Bieling et al., 2010; Funabiki and Murray, 2000; Takagi et al., 2013). hKid as well as XKid possesses an N-terminal motor domain, a short coiled-coil domain and a DNA binding tail domain (Figure 1A). Previous studies have shown that hKid and XKid, that are fused with fluorescent proteins at the C-terminal, can complement the function of hKid-depleted cells and XKid-depleted Xenopus egg extracts (Bieling et al., 2010; Soeda et al., 2016). Thus, we fused a fluorescent protein at the C-terminal of XKid and hKid. Notably, while EGFP-fused and sfGFP-fused hKid were insoluble, mNeonGreen-fused hKid was recovered from soluble fractions. As a result, we succeeded in purifying both XKid-mScarlet and hKid-mNeonGreen proteins (Figure 1B, and supplemental Figure S1). Next, the motility of purified proteins were analyzed by single molecule motility assays using total internal reflection fluorescent microscopy (TIRF). We found that both XKid and hKid moved on microtubules processively (Figure 1C, D, Movie 1 and 2). The motility of single molecules along microtubules could be observed at 20 pM of Xkid and hKid, respectively. The average velocity of XKid and hKid were approximately 110 nm/sec (Figure 1E and Table 1), within the same range as the movement of chromosomes (Brouhard and Hunt, 2005). The run length of XKid and hKid were 1.8 ± 0.7 and 1.3 ± 0.5 µm, respectively (Figure 1F and Table 1). These data collectively suggest that full-length Kid is a processive motor protein.

**Table 1.**
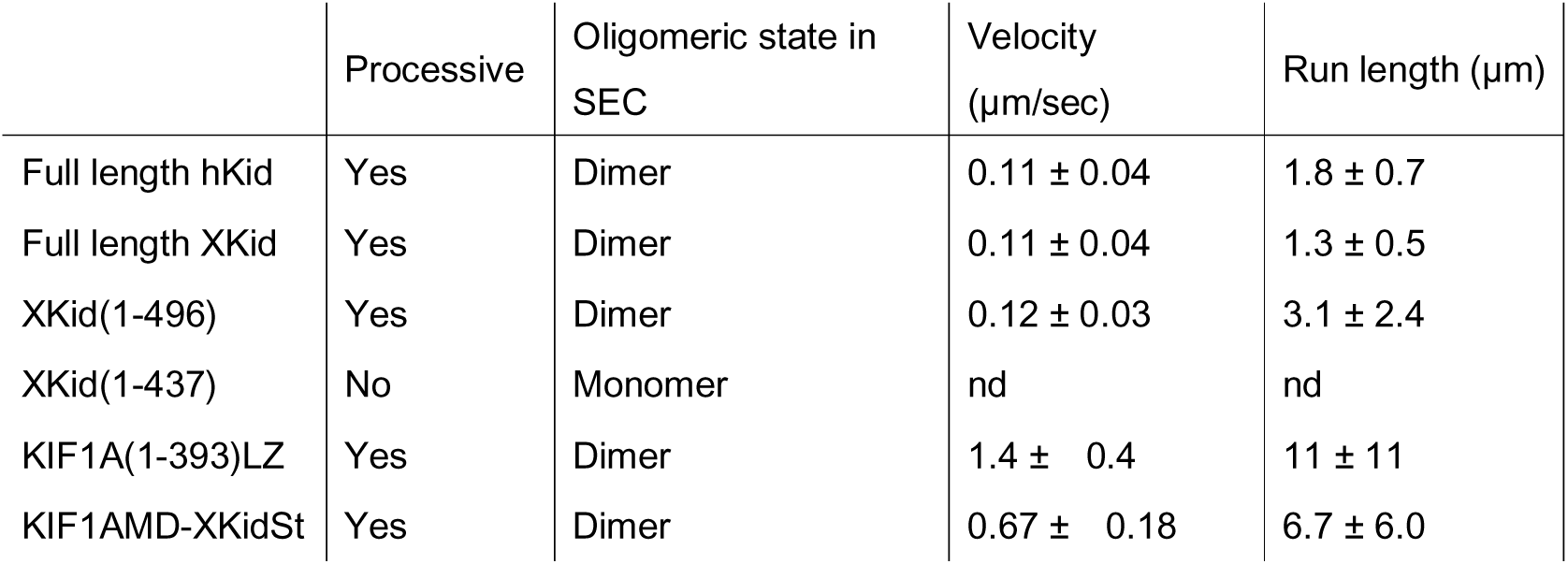
Motile properties of constructs used in this study. Motor protein constructs underwent purification through affinity chromatography followed by size exclusion chromatography, as detailed in the Materials and Methods section. The reported velocities and run lengths represent mean values ± standard deviation (SD). Notably, XKid(1-437) failed to demonstrate consistent processive motion across three separate protein preparations, indicating a lack of detectable activity (ND: not detected).

### Full-length Kid form dimers

It has been proposed that Kid is a monomeric protein (Shiroguchi et al., 2003; Yajima et al., 2003). Even full-length human Kid protein obtained from 293T cells are indicated to be a monomer (Shiroguchi et al., 2003). However, typical kinesins show processive movement along microtubules only when they form dimers (Hancock and Howard, 1998; Kita et al., 2024; Tomishige et al., 2002). As both hKid and XKid showed processive movement on microtubules (Fig 1), we next analyzed the oligomeric state of these motors using size exclusion chromatography (Fig 2A and B). We utilized the previously well-characterized kinesin UNC-104(1-653)-sfGFP as a size marker. UNC-104(1-653)-sfGFP exhibits a dimer peak at 200 kDa and a monomer peak at 100 kDa in size exclusion chromatography (Kita et al., 2024). This characteristic makes it a suitable marker for determining the oligomeric state of hKid and XKid, given that the predicted molecular weights of hKid-mNeonGreen and XKid-mScarlet monomers are approximately 100 kDa (Figure 1A). As a result, we found that the peak fraction of full-length hKid and XKid are almost equivalent to that of UNC-104(1-653)-sfGFP dimer (Fig 2A and B, and supplemental Figure S2). Notably, unlike UNC-104(1-653)-sfGFP, we did not detect any monomer peaks for hKid and XKid under these conditions. We next analyzed purified proteins recovered from the peak fractions using mass photometry (Sonn-Segev et al., 2020). Mass photometry is generally performed at the nanomolar concentrations (Sonn-Segev et al., 2020). Peak fractions obtained from size exclusion chromatography were diluted 100-fold and analyzed by mass photometry. Mass photometry analysis revealed a predominant monomer population, with a smaller fraction corresponding to dimers (Figure 2C and D). The behavior is similar to that of UNC-104(1-653)-sfGFP, which shows dimer peaks by size-exclusion chromatography at micromolar concentrations but dissociates into monomers at the nanomolar concentrations used for mass photometry (Kita et al., 2024). In addition, we consistently detected a minor trimer populations under this condition (Figure 2C and D), although this species may represent an artifact because it was not detected by size-exclusion chromatography. These findings indicate that full-length hKid and XKid are capable of forming dimers, but the dimer formation is dependent on the protein concentration.

**Figure 2.**
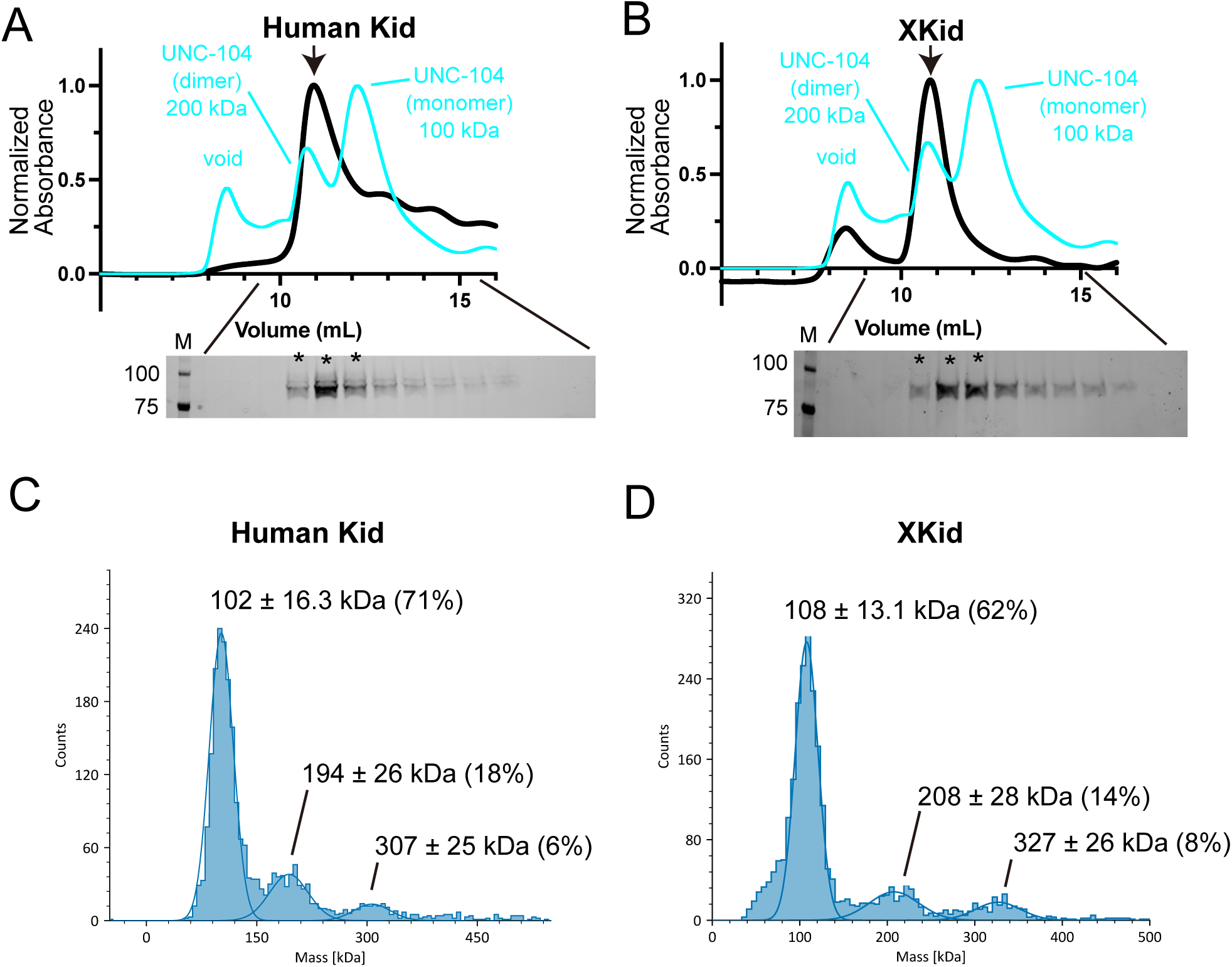
Kid forms a weak dimer. (A) Size exclusion chromatography profiles of hKidFL (black) and UNC-104(1-653)-sfGFP (cyan). Below the chromatography, an SDS-PAGE image show the elution fractions. Asterisks indicate fractions used for mass photometry and single molecule assays. The molecular weight standards are indicated on the left side of the SDS-PAGE images. (B) Size exclusion chromatography of XKidFL (black) and UNC-104(1-653) (cyan). The SDS-PAGE of the elution fractions are shown beneath the profiles. Asterisks indicate fractions used for mass photometry and single molecule assays. The number shown at the left side indicates molecular weight standard. (C) Mass photometry analysis of human Kid at 10 nM. Histograms show particle counts, and lines indicate Gaussian fits. The mean ± SD and percentage of each peak are shown. (D) Mass photometry analysis of human Kid at 10 nM. Histograms show particle counts, and lines indicate Gaussian fits. The mean ± SD and percentage of each peak are shown. Note that majority of hKid and XKid are dimers in the size exclusion chromatography but they are mostly dissociated to monomers in mass photometry.

### A conserved coiled-coil domain is essential for the processivity

To determine the domain essential for the processive movement of Kid, we generated a series of deletion mutants. Unfortunately, we failed to purify adequate quantities and qualities of hKid deletion mutants. Thus, following experiments were performed using XKid. We purified XKid(1-496), which lacks the DNA-binding tail domain, and XKid(1-437), which lacks the DNA-binding domain and the coiled-coil domain (Figure 3A and B, and supplemental Figure S1). Mass photometry showed that XKid(1-496) was predominantly monomeric (Fig 3C), with minor dimer and trimer populations, resembling the behavior of full length XKid (Figure 2). In contrast, XKid(1-437) was exclusively monomeric (Fig 3D).

**Figure 3.**
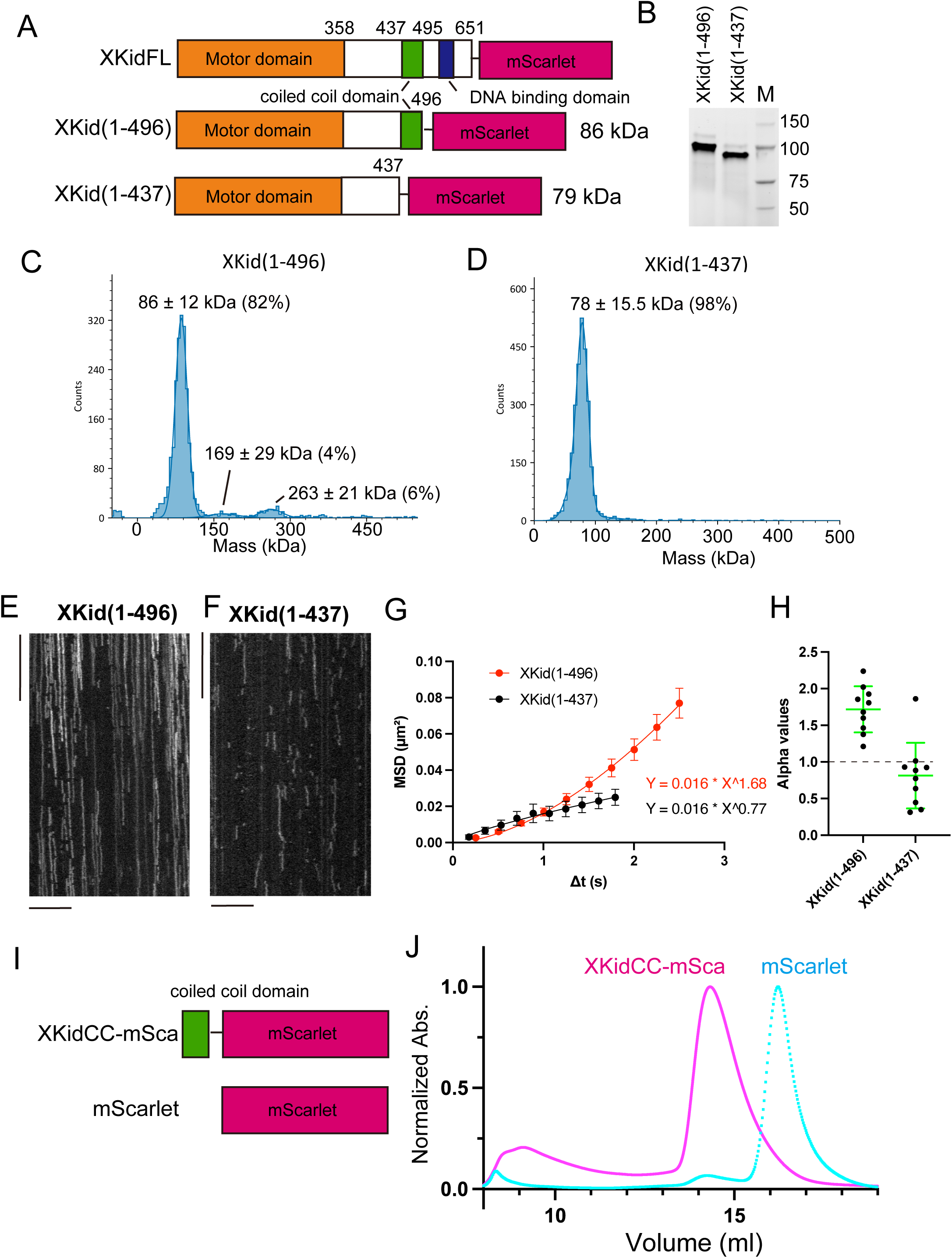
Conserved coiled-coil domain is required for the processive motion. (A) Schematic representation illustrating the domain organization of XKid(1-496) and XKid(1-437). The calculated molecular weights of the fusion proteins are indicated on the right. (B) Representative SDS-PAGE analysis of purified XKid(1-496) and XKid(1-437) proteins. The proteins are visualized using a Stain-Free gel. The molecular weight standards are indicated on the right side of the SDS-PAGE images. (C and D) Mass photometry analysis of XKid(1–496)-mSca and XKid(1–437)-mSca. The expected molecular masses are 86 and 79 kDa, respectively. Histograms show particle counts, and lines indicate Gaussian fits. The mean ± SD and percentage of total counts for each peak are shown. (E and F) Representative kymographs showing the motility of 10 pM XKid(1-496) (E) and XKid(1-437) (F) in the presence of 2 mM ATP. Note that no directional movement was detected in XKid(1-437). Scale bars: horizontal 10 µm; vertical 10 seconds. (G and H) Mean-square displacement (MSD) analysis of XKid(1–496) and XKid(1–437) trajectories. (G) Representative MSD curves fitted to the power-law relationship MSD = AΔt^α^, where α is the anomalous diffusion exponent. XKid(1–496) showed superlinear MSD scaling with α = 1.68, consistent with persistent or directionally biased motion, whereas XKid(1–437) showed sublinear MSD scaling with α = 0.77. (H) Distribution of α values obtained from individual trajectory fits. Each dot represents one trajectory; bars indicate mean ± SD. n = 10 trajectories per construct. (I and J) Schematic drawing of XKidCC-mScarlet (I) and a representative result of size exclusion chromatography (J). XKidCC-mScarlet (magenta) and mScarlet (cyan) are shown.

Using these purified proteins, we performed single molecule motility assays (Fig 3E-H). We found XKid(1-496) could move processively on microtubules (Fig 3E). The velocity of XKid(1-496) was approximately 120 nm/sec and the run length was 3.1 ± 2.4 µm, which are comparable to those of full-length XKid (Table 1). Thus, DNA binding domain of Kid is not required for processive runs. In contrast, XKid(1-437), which lacks the coiled-coil domain, did not show any processive runs (Fig 3F). To analyze the motile properties, we performed the mean square displacement (MSD) analysis (Fig 3 G and H). The MSD curves were fitted to the power-law relationship MSD = A(Δt)^α^, where α describes how the displacement scales with time. α = 1 indicates simple diffusion, α > 1 indicates superlinear MSD scaling indicating persistent or directionally biased motion, and α < 1 indicates sublinear MSD scaling indicating constrained motion. XKid(1–496) had an α value of approximately 1.6, consistent with directionally biased movement, whereas XKid(1–437) had an α value of approximately 0.8, indicating sublinear MSD scaling consistent with hindered motion (Figs. 3G 3H). This would be because XKid(1-437) did not form a homodimer, considering that coiled-coil domains are generally required for the dimerization of kinesins (Hancock and Howard, 1998; Kita et al., 2024).

To confirm that the coiled-coil domain of XKid induces dimerization, the domain was fused with mScarlet and analyzed by the size exclusion chromatography (Fig 3I and J). As a result, the peak of XKidCC-mScarlet shifted to the larger size compared with that of mScarlet. The calculated molecular weight of XKidCC-mScarlet was 42 kDa, which is almost equivalent to the size of dimerized mScarlet. To test whether monomeric Kid molecules in solution form dimers on microtubules, hKid-mScarlet3 and hKid-mStayGold were purified separately, mixed at final concentrations of 1 pM each, and analyzed using single-molecule motility assays (Fig 4). Under these assay conditions, hKid is expected to be monomeric in solution, because mass photometry showed that hKid was predominantly monomeric even at a much higher concentration of 10 nM (Figure 2). As a result, approximately 20% of motile hKid-mScarlet3 particles showed comigration with hKid-mStayGold particles along microtubules, suggesting microtubule-dependent association between hKid molecules.

**Figure 4.**
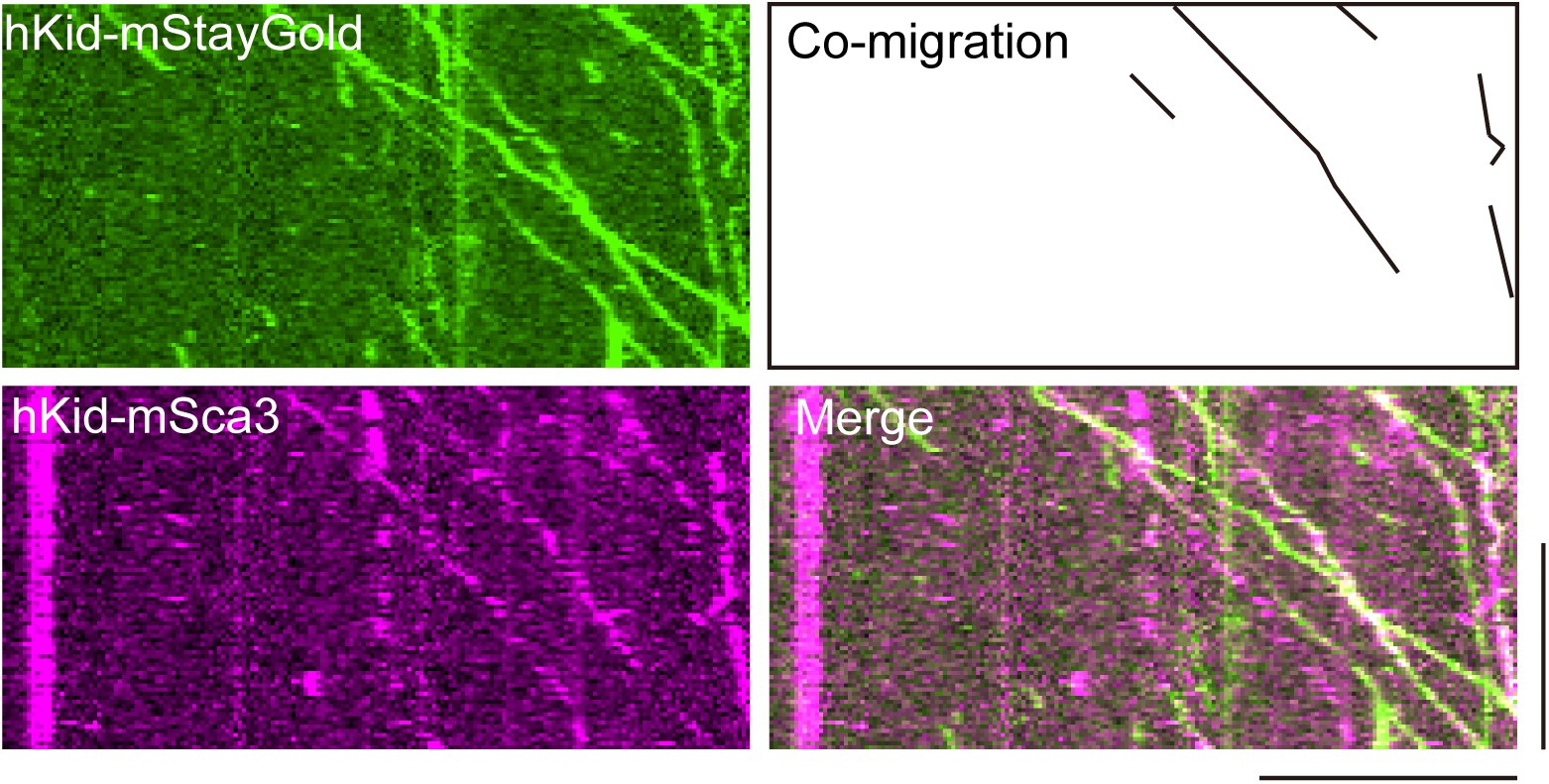
Human Kid–mStayGold and Kid–mScarlet3 co-migrate on microtubules. Purified hKid–mStayGold and hKid–mScarlet3 were mixed at final concentrations of 20 pM each and analyzed by single-molecule motility assays using TIRF microscopy. Representative kymographs show hKid–mStayGold, hKid–mScarlet3, and merged signals. The schematic drawing illustrates examples of co-migrating particles on microtubules, defined as overlapping mStayGold and mScarlet3 signals moving together along the same microtubule. Co-migration was defined as overlapping mStayGold and mScarlet3 signals moving together along the same microtubule over the same time interval. Scale bars: horizontal, 10 µm; vertical, 100 s.

### The stalk domain of XKid adds processivity to the motor domain of KIF1A

In the processive movement along the microtubules, the coordination between two motor domains is essential (Hancock and Howard, 1998). The neck linker domain, immediately following the kinesin motor domain, regulates the ATPase activity within the motor domain (Case et al., 2000). It has been shown that optimal length of the neck linker domain is crucial for achieving coordination of the two motor domains (Isojima et al., 2010; Yildiz et al., 2008). The length of the neck linker domain is typically determined by the presence of the neck coiled-coil domain (Case et al., 2000; Shastry and Hancock, 2010). However, Kid is an exception, as it does not have the conventional neck coiled-coil domain (Shiroguchi et al., 2003; Tokai et al., 1996). This unique feature of Kid supports the idea that Kid functions as a non-processive monomer (Yajima et al., 2003), which does not require the coordination between two motor domains. However, coiled-coil prediction tools might not identify hidden coiled-coil domains or Kid could have a motif that induces dimerization. In the case of kinesin-1, Alphafold2, but not coiled-coil prediction tools, can more accurately find the location of coiled-coil domains (Tan et al., 2023; Weijman et al., 2022). Therefore, we used Alphafold2 to analyze the neck-linker and the first coiled-coil domain of XKid and hKid. The regions corresponding to residues 359–495 of XKid and 369–506 of hKid were modeled using AlphaFold 2 (Figure 1A). The result suggested that the region does not have hidden coiled-coil domains nor a motif that induces dimerization, and the region is flexible (Supplementary Figure S3). If the entire flexible region functions as a neck linker, its length is 4 times longer than that of kinesin-1 (Supplementary Figure S3). To investigate if this extended neck linker of Kid can support the coordination of two motor domains, we fused the coiled-coil domain of XKid to the motor domain of KIF1A (Figure 5A). We included the possible neck linker domain of XKid in this chimera protein to test whether the neck linker of XKid is functional or not (Figure 5B). We could purify the chimeric protein, named KIF1AMD-XKidSt (Figure 5C and supplementary Figure S1). We found that KIF1AMD-XKidSt exhibited processive movement along microtubules in the single molecule motility assay (Figure 5D and movie S3), as is the case of KIF1A(1-393)LZ (Figure 5E and movie S4). The velocity of KIF1AMD-XKidSt was much faster than original XKid but slightly slower than KIF1A(1-393)LZ (Figure 5F and Table 1). The run length of KIF1AMD-XKidSt was shorter than KIF1A(1-393)LZ (Figure 5G and Table 1). Previous studies have shown that the motor domain of KIF1A does not exhibit processive motion when it is monomeric, but exhibits processive motion when an artificial dimer is generated using a stalk domain of kinesin-1 or a leucine zipper domain (Soppina et al., 2014; Tomishige et al., 2002). Therefore, these results suggest that the coiled-coil domain of XKid can induce dimerization of KIF1A motor domains on microtubules and the longer neck linker domain of XKid can support the processive movement of the kinesin motor domain.

**Figure 5.**
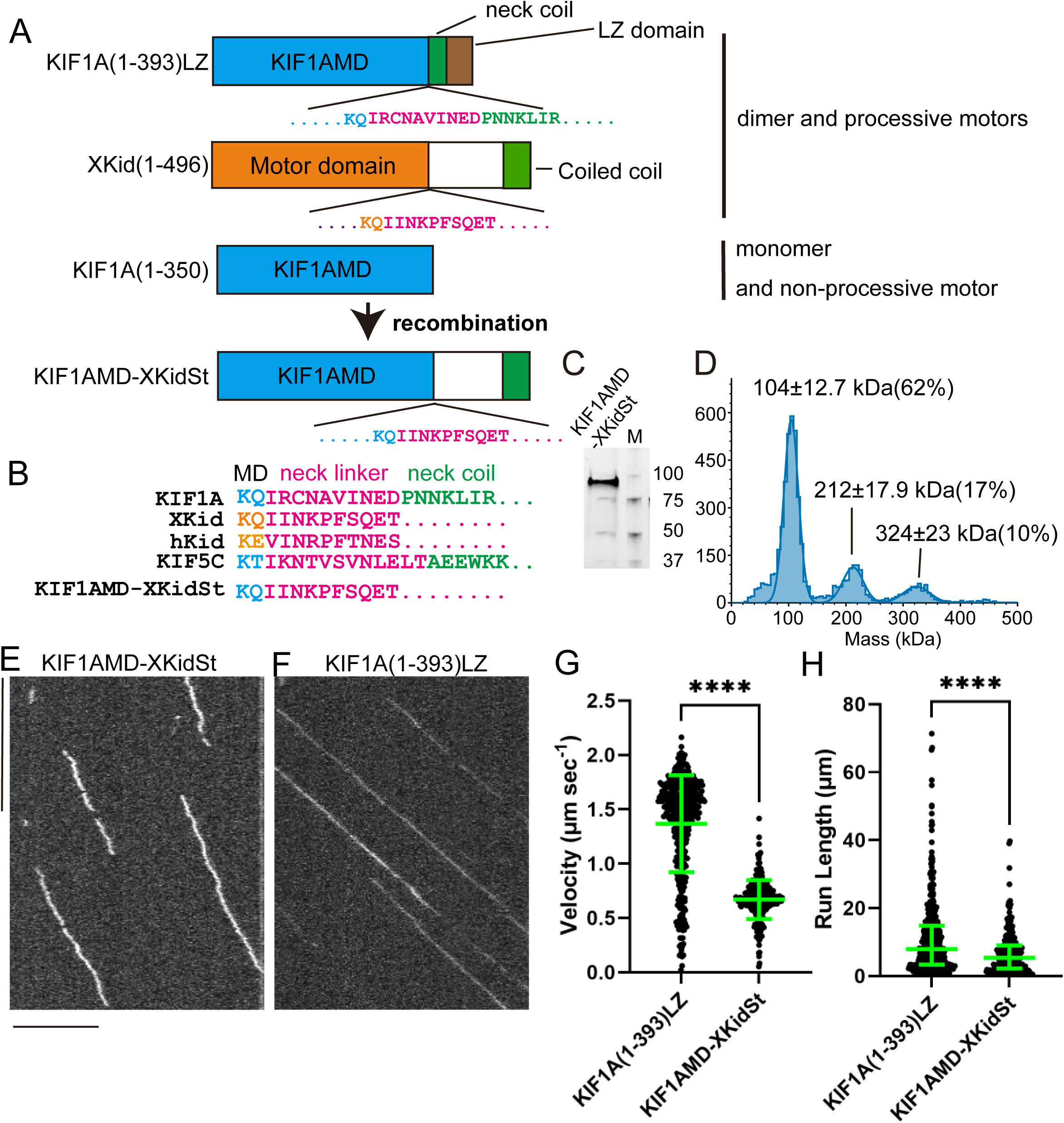
Untypical neck linker of Kid can support processive movement of KIF1A. (A) Schematic representation illustrating the domain organization of KIF1A(1-393)LZ, XKid(1-496), KIF1A(1-350) and a chimera protein KIF1AMD-XKidSt. Note that KIF1A(1-393)LZ and XKid(1-496) are processive motors and KIF1A(1-350) is a non-processive motor. Cyan, motor domain of KIF1A; Orange, motor domain of Kid; Magenta, neck linker. (B) Amino acid sequences of the neck linker region. KIF1A, XKid, hKid, KIF5C and KIF1AMD-XKidSt are shown. Cyan, motor domain of KIF1A and KIF5C; Orange, motor domain of Kid; Magenta, neck linker; Green, neck coiled-coil domain. (C) Representative SDS-PAGE analysis of purified KIF1AMD-XKidSt fusion protein. The protein is visualized using a Stain-Free gel. The molecular weight standards are indicated on the right side of the SDS-PAGE image. (D and E) Representative kymographs showing the motility of KIF1AMD-XKidSt and KIF1A(1-393)LZ in the presence of 2 mM ATP. Note that KIF1AMD-XKidSt exhibits diffusion-like fluctuations while they are moving. This phenomena is not observed in KIF1A(1-393)LZ. Scale bars: horizontal 10 µm; vertical 10 seconds. (F) Dot plots showing the velocity of KIF1AMD-XKidSt and KIF1A(1-393)LZ. Each dot shows a single datum point. Green bars represent mean ± S.D.. ****, p < 0.0001, Unpaired t-test. n = 273 and 434 particles for KIF1AMD-XKidSt and KIF1A(1-393)LZ, respectively. (G) Dot plots showing the run length of KIF1AMD-XKidSt and KIF1A(1-393)LZ. Each dot shows a single datum point. Green bars represent median value and interquartile range. ****, p < 0.0001, Mann-Whitney test. n = 273 and 434 particles for KIF1AMD-XKidSt and KIF1A(1-393)LZ, respectively.

### Reconstitution of DNA transport in vitro

Kid was originally identified as a DNA binding protein (Tokai et al., 1996). The tail domain of Kid has two helix-hairpin-helix motif which is supposed to be a DNA binding domain (Doherty et al., 1996). To test that Kid has an activity to transport DNA along microtubules, we mixed fluorescently-labelled DNA in TIRF assays and directly observed the motility (Figs 6A-D). To study the domains essential for the transport of DNA, we used full-length XKid and a deletion mutant of Xkid for these assays. We found that full-length XKid can drive the movement of double-stranded DNA along microtubules (Fig. 6A). Double-stranded DNA signals that co-migrated with XKid were 93.4 ± 10.5 % (n = 31 microtubules). The velocity of double-stranded DNA moving along microtubules was approximately 100 nm/sec, which is similar to the velocity of XKid alone (Figure 1). In contrast, single-strand DNA movement was not driven by XKid (Fig. 6B and D). Deletion of the tail domain of XKid, containing helix-hairpin-helix motif, abolished the DNA transport activity (Fig 6C and D). We confirmed that full-length hKid can also induce the movement of double-stranded DNA along microtubules (Figure S4 and supplemental movie S5).

**Figure 6.**
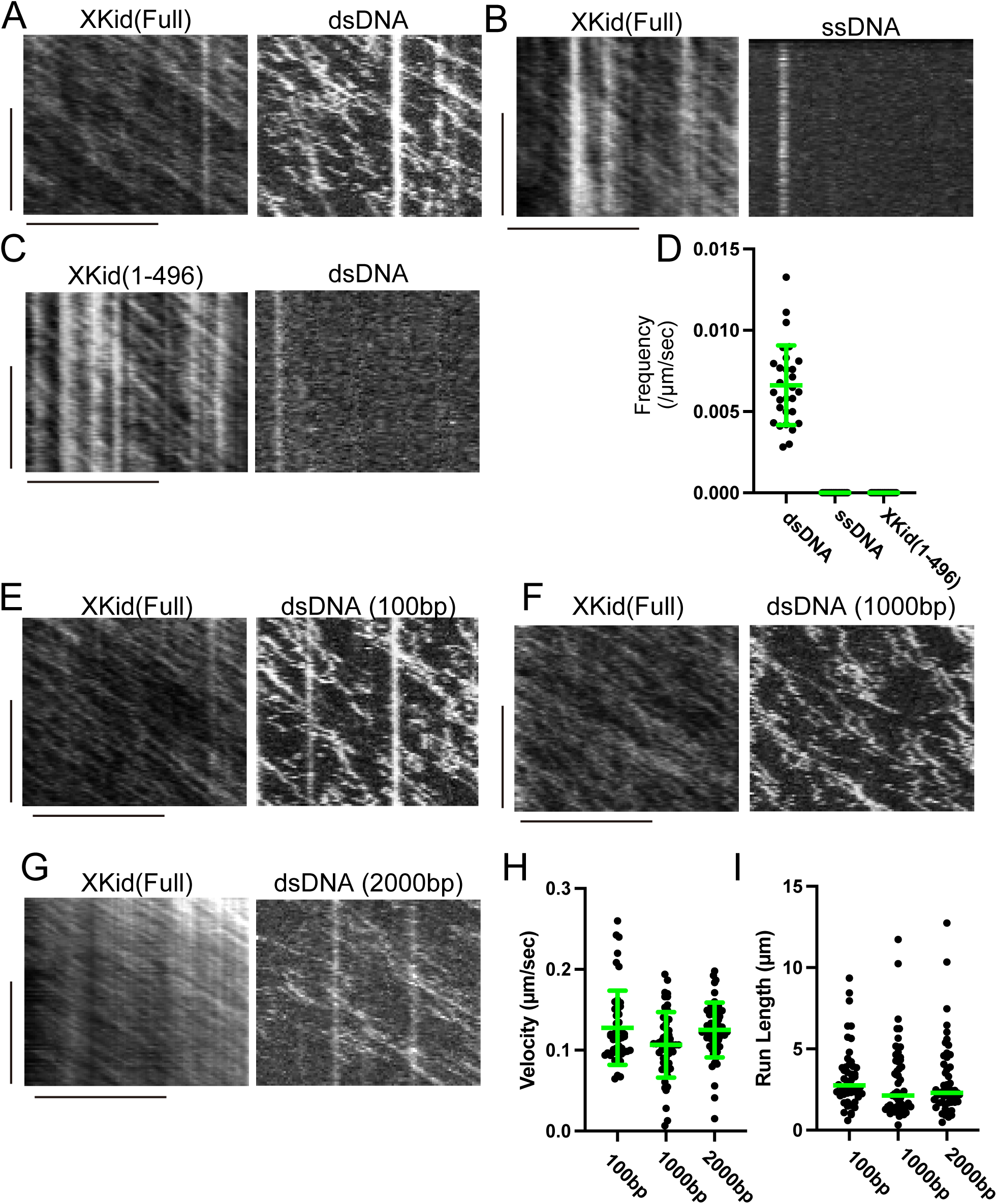
DNA movement driven by XKid. (A–D) sfGFP-tagged XKid at 1 nM was mixed with 20 nM Cy3-labeled double-stranded or single-stranded DNA and observed by TIRF microscopy. (A–C) Representative kymographs showing the movement of full-length XKid with double-stranded 100-bp DNA (A), full-length XKid with single-stranded 100-base DNA (B), and XKid(1–496) with double-stranded 100-bp DNA (C). Scale bars: horizontal, 10 µm; vertical, 100 s. (D) Frequency of DNA movement along microtubules, normalized by microtubule length and observation time. Each dot represents an individual measurement from a different microtubule. Bars indicate mean ± SD. n = 29 microtubules per condition. (E–I) Single-molecule analysis of XKid–sfGFP on Cy3-labeled DNA. Purified XKid–sfGFP was mixed with Cy3-labeled DNA at final concentrations of 1 nM XKid–sfGFP and 20 nM DNA and observed by TIRF microscopy. (E–G) Representative kymographs showing the movement of full-length XKid–sfGFP on 100-bp DNA (E), 1000-bp DNA (13 ng/µl; F), and 2000-bp DNA (26 ng/µl; G). Scale bars: horizontal, 10 µm; vertical, 100 s. (H) Velocity of DNA movement along microtubules. Each dot represents one DNA molecule. Bars indicate mean ± SD. n = 48, 53, and 53 DNA molecules for 100-, 1000-, and 2000-bp DNA, respectively. (I) Run length of DNA movement along microtubules. Each dot represents one DNA molecule. Bars indicate mean ± SD. n = 48, 53, and 53 DNA molecules for 100-, 1000-, and 2000-bp DNA, respectively.

Next, we analyzed dsDNA fragments of different lengths. Full-length Kid transported not only 100-bp dsDNA but also longer 1,000-bp and 2,000-bp dsDNA fragments along microtubules (Fig. 6E-G). However, the velocity and run length of DNA transport were comparable among these substrates (Fig. 6H and I), suggesting that increasing DNA length does not substantially alter Kid-driven DNA transport under our reconstituted assay conditions.

### Dimerization would be required for DNA transport

The structure of the XKid–DNA complex was modeled using AlphaFold3 (Fig 7A-D). We first modeled one copy of the DNA-binding domain of XKid with a 15-bp DNA fragment (Fig 7A). This prediction yielded low ipTM and pTM values of 0.12 and 0.64, respectively, and did not produce a clear XKid-DNA complex model. We then modeled two copies of the DNA-binding domain of XKid with a double-stranded DNA fragment (Figure 7B–D). In this case, AlphaFold 3 generated a model in which two DNA-binding domain of XKid associate with dsDNA. The ipTM and pTM values were 0.79 and 0.84, respectively, suggesting a higher-confidence prediction of the complex. To further examine whether the DNA-binding domain of XKid intrinsically forms a stable dimer, we performed AlphaFold3 prediction in the absence of DNA. This prediction yielded only modest confidence scores (ipTM = 0.34, pTM = 0.57). These results suggest that the dimeric configuration of the DNA-binding domain of XKid is preferentially stabilized in the presence of double-stranded DNA. These results suggest that the DNA-binding domain of XKid adopts a DNA-binding architecture more readily in a dimeric configuration. To experimentally test the predicted model, we introduced the K613A mutation into XKid, as K613 was predicted to be located at the XKid–DNA interface (Fig. 7C), and performed dsDNA motility assays (Fig. 7D,E). The XKid(K613A) mutant failed to drive DNA movement in the reconstitution assay (Fig. 7E).

**Figure 7.**
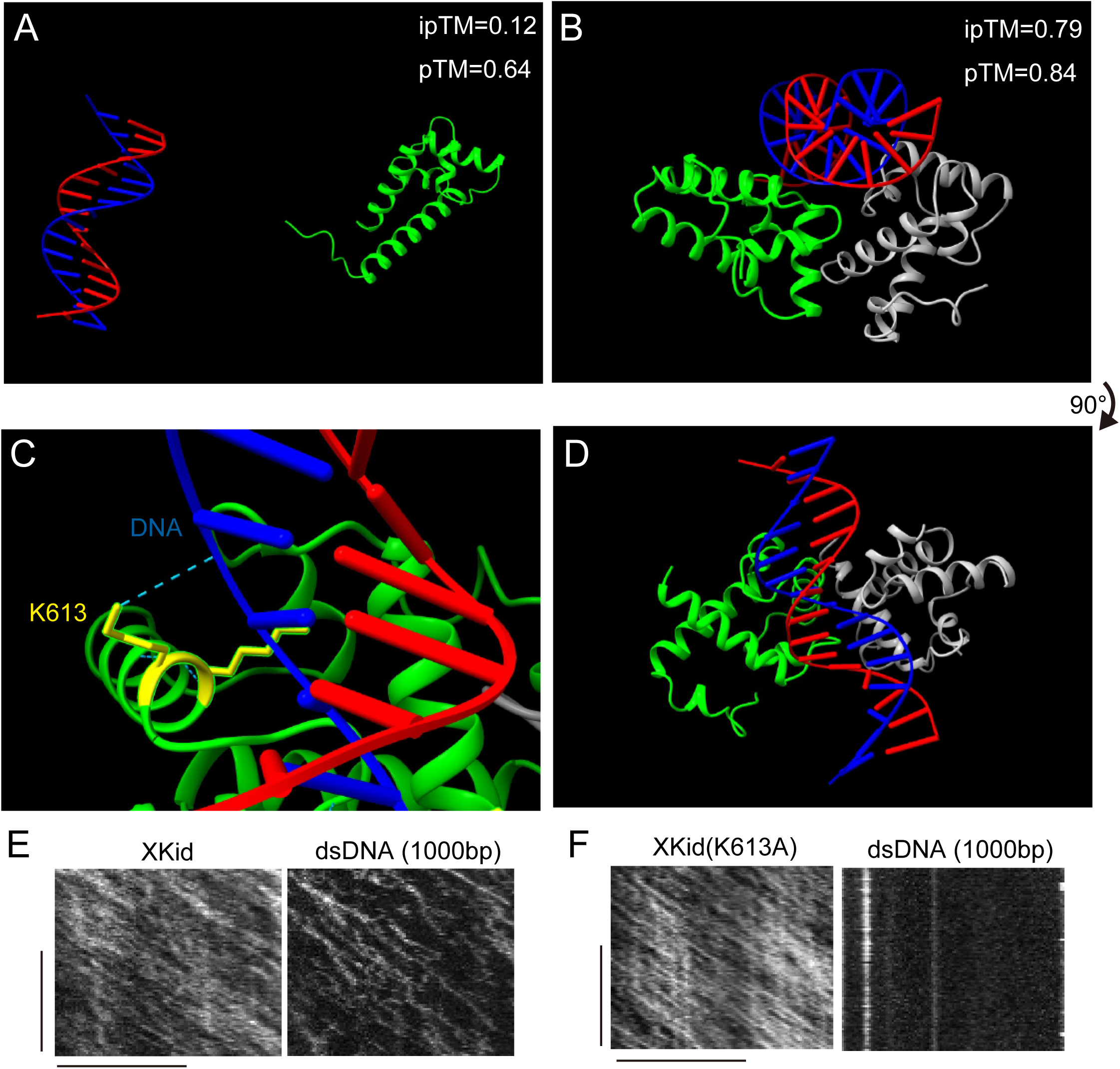
AlphaFold3 prediction and functional validation. (A) AlphaFold3-predicted structure of the XKid DNA-binding domain modeled as a monomer with double-stranded DNA (dsDNA). The prediction showed a low interface confidence score between XKid and dsDNA (ipTM = 0.12, pTM = 0.64). (B–D) AlphaFold3-predicted structure of the XKid DNA-binding domain modeled as a dimer with dsDNA. The two XKid molecules are shown in green and gray, and the two DNA strands are shown in red and blue. This dimeric model showed higher confidence for the XKid–DNA complex (ipTM = 0.79, pTM = 0.84). (B) Side view of the predicted XKid dimer–dsDNA complex. (C) Magnified view of the predicted XKid–DNA interface. K613 and K614, highlighted in yellow, are positioned near dsDNA. A cyan dashed line indicates the predicted hydrogen bond between K613 and DNA. (D) View of the same model shown in (B) after a 90° rotation. (E and F) Representative kymographs showing the movement of dsDNA along microtubules. Wild-type XKid supported movement of 1000-bp dsDNA (E), whereas the K613A mutant abolished detectable dsDNA movement (F). Scale bars: horizontal, 10 µm; vertical, 100 s.

## Discussion

### Kid is a processive dimer on microtubules

Prometaphase chromosomes are transported along microtubules by the activity of kinesins (Iemura and Tanaka, 2015; Wordeman, 2010). Kid is the primary kinesin that transports chromosomes along spindle microtubules in prometaphase (Brouhard and Hunt, 2005; Iemura and Tanaka, 2015). Previous biochemical studies reported that purified Kid is monomeric, leading to the concept that Kid is a monomeric and non-processive chromokinesin (Yajima et al., 2003; Shiroguchi et al., 2003). Under this model, sustained chromosome movement would require many Kid monomers distributed along chromosome arms to act collectively. Our findings revise this view. Our data show that full-length Kid is capable to form a dimer and exhibits a processive motion along microtubules. In addition, Kid dimers directly bind to dsDNA. Thus, the elementary force-generating unit of Kid is a single Kid dimer which functions as a processive DNA-bound motor. In the context of mitotic chromosomes, multiple processive Kid dimers bound along chromosome arms could cooperate to generate chromosome-scale polar ejection forces (Figure 8). This model preserves the likely importance of motor team on large chromatin. However, the team is likely composed of multiple processive dimers rather than many non-processive monomers.

**Figure 8.**
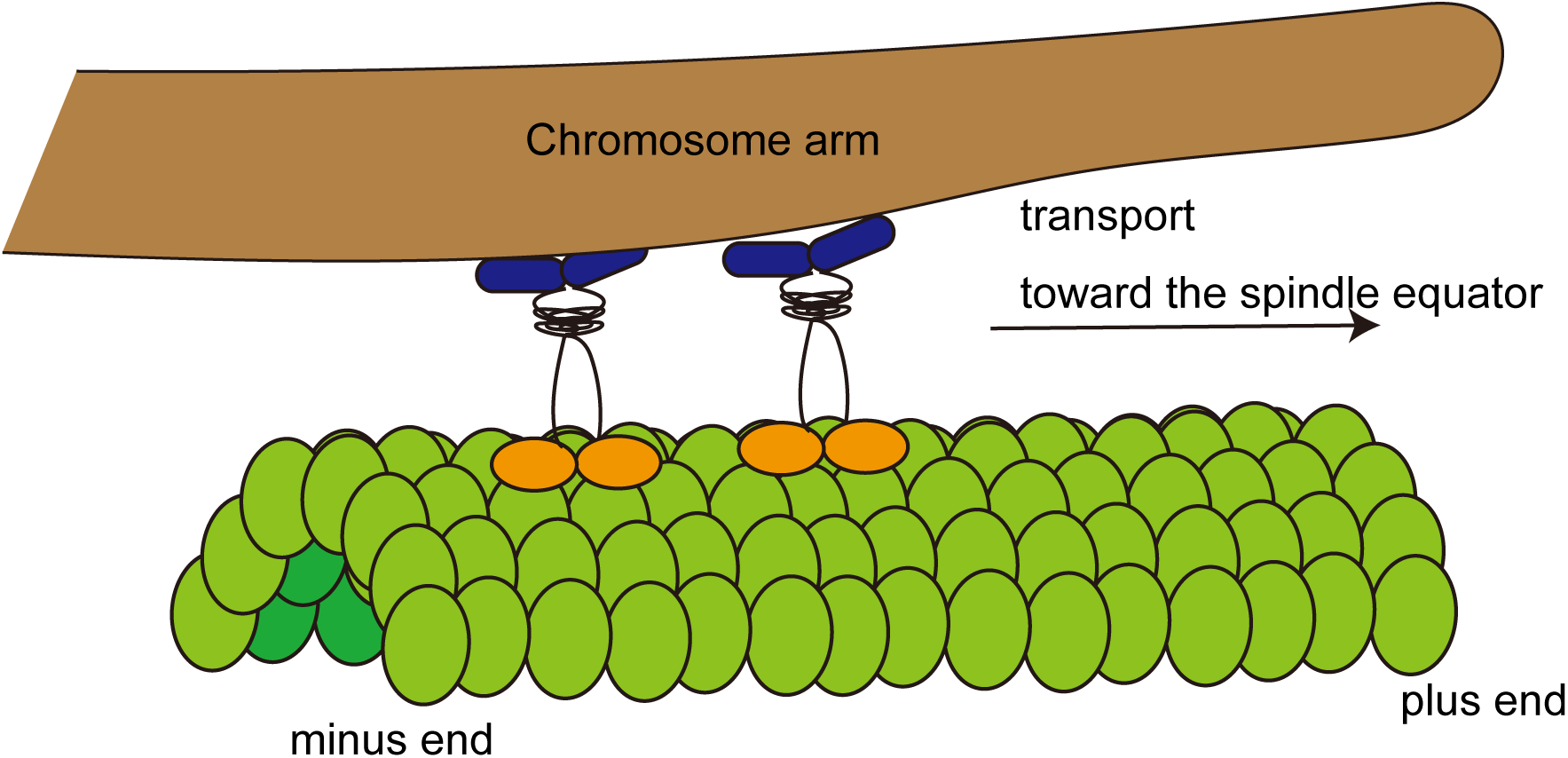
Model. Model for Kid-dependent DNA and chromosome transport. Kid forms processive dimers that can directly bind and transport double-stranded DNA along microtubules. On mitotic chromosomes, multiple Kid dimers may bind along chromosome arms and cooperate to generate polar ejection forces. This model is not drawn to scale and does not fully represent the structural complexity of condensed chromatin.

### Biochemical properties of Kid

A notable difference between this study and prior studies is the protein concentration used. Previous analysis of full-length human Kid via density gradient and size exclusion chromatography identified peak fractions through western blot, due to the low concentration of recombinant Kid expressed in 293T cells (Shiroguchi et al., 2003). We show that most hKid and XKid exist as dimers in size exclusion chromatography performed at micromolar concentrations, yet mostly dissociate into monomers in mass photometry, which is performed at nanomolar concentrations (Figure 2). Thus, the inability to detect Kid dimers in earlier studies may be attributed to the low concentration. In mass photometry, we also detected a small population with an apparent trimeric mass (Figs. 2, 3, and 5). Because this species was not detected by size-exclusion chromatography, we interpret this population cautiously and cannot exclude the possibility that it represents a measurement or sample-preparation artifact rather than a stable oligomeric state of Kid. However, transient higher-order oligomerization of Kid in cells, where Kid may be locally concentrated on microtubules or chromatin, remains possible and should be examined in future studies.

Another difference is the methods to detect the processivity of motors. We observed the processivity of Kid by TIRF whereas previous studies observed the motility by optical trap. In optical trapping assays, Kid is diluted prior to being adsorbed onto beads. This dilution likely promotes the dissociation of preformed Kid dimers, consistent with our mass photometry analysis showing that Kid is predominantly monomeric at nanomolar concentrations (Fig. 2), and thus optical trapping assays would primarily monitor monomeric Kid motility. By contrast, in TIRF-based motility assays, Kid in solution binds directly to microtubules. Although Kid is predominantly monomeric in solution, its accumulation on the microtubule surface may increase the local concentration of Kid and promote dimer formation on microtubules. This microtubule-dependent dimerization model could account for the processive motility observed in the TIRF assay. Consistent with this model, when separately purified Kid-mScarlet3 and Kid-mStayGold were mixed, they colocalized and moved processively together along microtubules (Figure 5). However, because of the limited temporal resolution of our two-color imaging system, we could not directly resolve the transition from monomers to dimers on microtubules. These properties, such as “the equilibrium between monomers and dimers” and “the capability to exhibit processive movement on microtubules, even when characterized as monomers through mass photometry” have been observed in kinesin-3 motors (Chiba et al., 2023; Fan, 2022; Kita et al., 2024).

### Neck linker of Kid

We show that Kid has a exceptionally long neck linker, approximately four times longer than that of kinesin-1. Neck-linker length strongly influences kinesin processivity, and changes in neck-linker length alter the run length and motility properties of kinesin-1, kinesin-2, and other N-terminal kinesins (Shastry and Hancock, 2010; Shastry and Hancock, 2011). However, longer or non-conventional neck linker regions can also support processive motility and may provide additional functions, such as navigation around microtubule-bound obstacles. Kinesin-2 bypasses Tau and other microtubule-bound obstacles by protofilament switching (Hoeprich et al., 2014). The neck linker of the mitotic kinesin KIF18A contributes to obstacle navigation within the mitotic spindle (Malaby et al., 2019). Thus, the exceptionally long and flexible neck linker of Kid may represent an adaptation that allows this chromokinesin to move processively along crowded spindle microtubules while remaining attached to DNA or chromatin. This possibility remains to be tested directly.

### Reconstitution of chromosome congression

A previous in vitro study, using an elegant assay termed chromatin gliding assay, has shown that XKid and XKLP1/KIF4A can crosslink between DNA and microtubules (Bieling et al., 2010). However, it remains to be elusive whether the chromosome transport is mediated by direct binding between chromosomal DNA and motor proteins. In organelle transport, cargo adaptor proteins are generally required for efficient transport (Chiba and Niwa, 2024; Chiba et al., 2022). Our in vitro reconstitution suggests that Kid-dependent chromosome transport does not require cargo adaptor proteins. Rather, our results suggest that dimerization of Kid facilitates DNA binding, allowing Kid to directly couple DNA to microtubule-based motility (Figure 7). Although hKid transported 1,000-bp and 2,000-bp DNA fragments in vitro, their motile parameters were comparable to those of 100-bp DNA. Thus, increasing DNA length did not substantially enhance DNA transport under our reconstituted assay conditions. One possible explanation is that the interaction between Kid and naked DNA is relatively weak or dynamic, and thus only one or a small number of Kid molecules productively engage each DNA molecule during transport. Alternatively, additional Kid molecules bound to longer DNA may not strongly affect the measured motility parameters under these assay conditions. However, naked DNA does not fully recapitulate the structural and mechanical properties of condensed chromatin or mitotic chromosomes. Thus, although multiple Kid dimers may engage chromatin to generate chromosome-scale polar ejection forces, this model remains to be directly tested. Future experiments using chromatinized DNA or reconstituted chromosome-like substrates will be required to determine how Kid interacts with condensed chromatin and how multiple Kid molecules cooperate to move chromosomes during prometaphase.

It has been shown that Kid is regulated by phosphorylation by CDK1(Ohsugi et al., 2003). It would be interesting to study the effects of phosphorylation and dephosphorylation on the Kid-dependent DNA transport using this reconstitution system. Moreover, by extending this system, it may be possible to fully reconstitute the chromosome congression in vitro by including other kinesins and microtubule-associated proteins, such as XKLP1/KIF4A, CENP-E and NuSAP1, that are required for proper chromosome congression (Bieling et al., 2010; Iemura and Tanaka, 2015; Li et al., 2016).

## Methods

### Plasmids

PCR was performed using a KOD FX neo DNA polymerase (TOYOBO, Tokyo, Japan). Human Kid cDNA (corresponding to DQ895829.2) was described previously (Iemura and Tanaka, 2015). Xenopus Kid (Kif22.S, corresponding to BC070549.1) was purchased from Horizon Discovery. To generate hKidFL-mNeonGreen, DNA fragments encoding human Kid and mNeonGreen were amplified by PCR and assembled into pFastbac1 (Novagen) by Gibson assembly as described (Gibson et al., 2009). To generate XKidFL-mScarlet, DNA fragments encoding XKid and mScarlet were amplified by PCR and assembled into pAcebac1 (Geneva Biotech). ORF sequences are shown in Supplementary Table S1. Deletion mutants of XKid were generated by PCR-based mutagenesis. For this purpose, primers were designed through the QuickChange Primer Design tool, a web-based application (Agilent). PCR-based mutagenesis was performed using KOD plus neo DNA polymerase (TOYOBO).

### Expression of XKid and Kid in Sf9 cells

Sf9 cells (Thermo Fisher Scientific) were maintained in Sf900^TM^ II SFM (Thermo Fisher Scientific) at 27°C. DH10Bac (Thermo Fisher Scientific) were transformed to generate bacmid. To prepare baculovirus, 1 × 10^6^ cells of Sf9 cells were transferred to each well of a tissue-culture treated 6 well plate. After the cells attached to the bottom of the dishes, about ∼5 μg of bacmid were transfected using 5 μL of TransIT^®^-Insect transfection reagent (Takara Bio Inc.). 5 days after initial transfection, the culture media were collected and spun at 3,000 × g for 3 min to obtain the supernatant (P1). For protein expression, 400 mL of Sf9 cells (2 × 10^6^ cells/mL) were infected with 200 µL of P1 virus and cultured for 65 h at 27°C. Cells were harvested and stocked at -80°C.

### Purification of proteins

We failed to purify hKid-EGFP and hKid-superfolder GFP due to the insolubility. In contrast, mNeonGreen fusion and mStayGold fusion stabilized hKid and enabled purification.

Sf9 cells were resuspended in 40 mL of Kid lysis buffer (50 mM HEPES-KOH, pH 7.5, 500 mM KCH3COO, 2 mM MgSO4, 1 mM EGTA, 10% glycerol) along with 1 mM DTT, 1 mM PMSF, 0.1 mM ATP and 0.5% Triton X-100. After incubating on ice for 10 min, lysates were cleared by centrifugation (100,000 × g, 20 min, 4°C) and subjected to affinity chromatography. Lysate was loaded on Streptactin-XT resin (IBA Lifesciences, Göttingen, Germany) (bead volume: 2 ml). The resin was washed with 40 ml Kid wash buffer (50 mM HEPES-KOH, pH 8.0, 500 mM KCH_3_COO, 2 mM MgSO_4_, 1 mM EGTA, 10% glycerol). Protein was eluted with 40 ml Kid elution buffer (50 mM HEPES-KOH, pH 8.0, 500 mM KCH_3_COO, 2 mM MgSO_4_, 1 mM EGTA, 10% glycerol, 200 mM biotin). Eluted solution was concentrated using an Amicon Ultra 15 (Merck) and then separated on an NGC chromatography system (Bio-Rad) equipped with a Superdex 200 Increase 10/300 GL column (Cytiva). Peak fractions were collected and concentrated using an Amicon Ultra 4 (Merck). Proteins were analyzed by SDS-PAGE using TGX Stain-Free gel (Bio-Rad). Concentrated proteins were aliquoted and snap-frozen in liquid nitrogen.

### Mass photometry

Purified hKid and XKid obtained from the peak fractions in the SEC analysis were pooled, snap-frozen and stored until measurement. Prior to measurement, the proteins were thawed and diluted to a final concentration 5 - 10 nM in GF150 buffer (25 mM HEPES, 150 mM KCl, 2 mM MgCl_2_, pH 7.2). Mass photometry was performed using a Refeyn OneMP mass photometer (Refeyn) and Refeyn AcquireMP version 2.3 software, with default parameters set by Refeyn AcquireMP. Bovine serum albumin (BSA) was used as a control to determine the molecular weight. The results were subsequently analyzed and graphs were prepared to visualize the data using Refeyn DiscoverMP version 2.3.

### Preparation of microtubules

Tubulin was purified from porcine brain as described (Castoldi and Popov, 2003). Tubulin was labeled with Biotin-PEG_2_-NHS ester (Tokyo Chemical Industry, Tokyo, Japan) and AZDye647 NHS ester (Fluoroprobes, Scottsdale, AZ, USA) as described (Al-Bassam, 2014). To polymerize Taxol-stabilized microtubules labeled with biotin and AZDye647, 30 μM unlabeled tubulin, 1.5 μM biotin-labeled tubulin and 1.5 μM AZDye647-labeled tubulin were mixed in BRB80 buffer supplemented with 1 mM GTP and incubated for 15 min at 37°C. Then, an equal amount of BRB80 supplemented with 40 μM taxol was added and further incubated for more than 15 min. The solution was loaded on BRB80 supplemented with 300 mM sucrose and 20 μM taxol and ultracentrifuged at 100,000 g for 5 min at 30°C. The pellet was resuspended in BRB80 supplemented with 20 μM taxol.

### Preparation of fluorescent-labelled DNA for the TIRF assay

To prepare 100 bp DNA fragment, oligonucleotides labelled with Cy3 were purchased from Integrated DNA Technologies, Inc. (Coralville, Iowa, USA). Following oligonucleotides were synthesized; Cy3-5’-GAGAATCGCCGGTTGATAATCTT<u>CCTAGTAGGTAGTATTGGTGTTGAGTCGCTCA</u>-3’ (Oligonucleotide #1) Cy3-5’-GAGAATCGCCGGTTGATAATCTT<u>TGAGCGACTCAACACCAATACTACCTACTAGG</u>- 3’ (Oligonucleotide #2) Underlines indicate sequences that form double-stranded DNA.

Double-stranded DNA was prepared using either a VeritiPro Thermal Cycler (Applied Biosystems) or a C1000 Touch Thermal Cycler (Bio-Rad). 1 µM of Oligonucleotides #1 and #2 were mixed and subjected to the following protocol: an initial incubation at 96 °C for 2 minutes, followed by incubation at 25 °C for 1 minutes, and subsequently to 4 °C. For single-stranded DNA preparation, Oligonucleotide #1 was incubated at 96 °C and then cooled to 4 °C.

To prepare 1000-bp and 2000-bp DNAfragments, Cy3-labeled oligonucleotides were purchased from Eurofins Genomics K.K. (Tokyo, Japan). The following oligonucleotides were synthesized: Cy3-5’-TGATGACGGTGAAAACCTCTGACAC-3’ (pUC19_F) 5’-TATGAGAAAGCGCCACGCTTCCCG-3’ (pUC19_R_1000bp) 5’-GATCGGAGGACCGAAGGAGCTAACC-3’(pUC19_R_2000bp) PCR was performed using pUC19 as the template and KOD FX Neo DNA polymerase. After agarose gel electrophoresis, the 1,000-bp and 2,000-bp DNA fragments were purified.

Double-stranded and single-strand DNA were prepared on the day of the experiment, immediately before the TIRF single-molecule motility assays. Old DNA can potentially cause high background.

### TIRF single-molecule motility assays

Purified Kid proteins described above were thawed and analyzed. KIF1A (1-393)LZ, that was described in our previous work (Anazawa et al., 2022), was also thawed and reanalyzed. TIRF assays using porcine microtubules were performed as described (Chiba et al., 2019). Glass chambers were prepared by acid washing as previously described (Chiba et al., 2022). Glass chambers were coated with PLL-PEG-biotin (50% labelled, SuSoS, Dübendorf, Switzerland) and streptavidin (Wako). Polymerized microtubules were flowed into flow chambers and allowed to adhere for 5–10 min. Unbound microtubules were washed away using assay buffer (90 mM HEPES-KOH pH 7.4, 50 mM KCH_3_COO, 2 mM Mg(CH_3_COO)_2_, 1 mM EGTA, 10% glycerol, 0.1 mg/ml biotin–BSA, 0.2 mg/ml kappa-casein, 0.5% Pluronic F127, 2 mM ATP, and an oxygen scavenging system composed of PCA/PCD/Trolox). Purified Kid was diluted to indicated concentrations in the assay buffer. Then, the solution was flowed into the glass chamber. An ECLIPSE Ti2-E microscope equipped with a CFI Apochromat TIRF 100XC Oil objective lens (1.49 NA), an Andor iXion life 897 camera and a Ti2-LAPP illumination system (Nikon, Tokyo, Japan) was used to observe the motility. NIS-Elements AR software ver. 5.2 (Nikon) was used to control the system. At least three independent experiments were conducted for each measurement.

### MSD analysis

Single-particle trajectories of fluorescent XKid(1-496) and XKid(1-437) molecules were obtained by manually tracking fluorescent puncta in kymographs. For each trajectory, the MSD was calculated for each lag time, Δt, as:

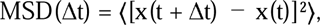

where x(t) is the one-dimensional position of the particle along the microtubule or DNA axis, and the brackets denote averaging over all displacement pairs with the same lag time. When the time intervals between tracked points were not strictly uniform, Δt was defined as the mean time difference of all displacement pairs contributing to that lag. The standard error of the MSD for each lag time was calculated as SD/√N, where N is the number of displacement pairs. To minimize the influence of reduced sampling at longer lag times and end-point effects, curve fitting was restricted to the initial region of each MSD curve. Unless otherwise indicated, lag times within approximately the first 60% of the total trajectory duration were used. Lag times were included in the fitting only when they were calculated from at least 10 displacement pairs. Each MSD curve was fitted to the power-law relationship:

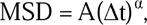

where A is a scaling coefficient and α is the anomalous diffusion exponent.

### Alphafold2 analysis

Alphafold2 analysis was conducted on Google Colaboratory (Jumper et al., 2021; Mirdita et al., 2022). We analyzed the amino acid sequences of KIF5C, XKid, and hKid, ranging from the neck linker to the end of coiled-coil 1. The sequences are detailed in Supplementary Table S2. The analysis was based on the assumption that these fragments form dimers and XKid and hKid fold in the similar manner.

### Alphafold3 analysis

Alphafold3 analysis was conducted on Alphafold Server (https://alphafoldserver.com/). We analyzed the amino acid sequences of the DNA binding domain of XKid and 15-bp dsDNA. The sequences are detailed in Supplementary Table S3.

### Statistical analyses and graph preparation

Statistical analyses and graph preparation were conducted using GraphPad Prism version 10. Details on the statistical methods are provided in the figure legends. Graphs were created with GraphPad Prism version 10, exported in PDF format, and aligned using Adobe Illustrator 2023.

## Acknowledgments

We would like to thank the members of Niwa lab for useful discussions. We also would like to thank Dr. Atsushi Nakagawa and Mr. Jiye Wang (Osaka University) for technical assistance. SN was supported by JSPS KAKENHI (grant no. JP23H02472). TK was supported by JSPS KAKENHI (grant no. JP23KJ0168). KC was supported by JSPS KAKENHI (grant no. JP22K15053), Uehara Memorial Foundation, Naito Foundation and MEXT Leading Initiative for Excellent Researchers (grant no. JPMXS0320200156). This work was performed under the Collaborative Research Program of Institute for Protein Research, Osaka University, CR-24-02.

## Author contributions

**Shinsuke Niwa:** Conceptualization; Resources; Data curation; Formal analysis; Project administration; Supervision; Funding acquisition; Validation; Investigation; Visualization; Methodology; Writing—original draft; Writing—review and editing.

**Natsuki Furusaki:** Resources; Data curation; Formal analysis; Validation; Investigation

**Tomoki Kita:** Resources; Data curation; Formal analysis; Validation; Investigation

**Kyoko Chiba:** Resources; Data curation; Formal analysis; Supervision; Funding acquisition; Project administration; Writing—review and editing.

## Statement

During the preparation of this work the authors used GPT4.0 in order to check English grammar and improve English writing. After using this tool, the authors reviewed and edited the content as needed and take full responsibility for the content of the publication.

## Supplemental Movie legends

### Movie S1

The motility of XKid was analyzed through time-lapse imaging recorded at 0.5 frames per second (fps) and played back at 25 fps. The dimensions of the displayed frame are 30 µm by 7 µm.

### Movie S2

The motility of hKid was analyzed through time-lapse imaging recorded at 0.5 fps and played back at 25 fps. The dimensions of the displayed frame are 30 µm by 7 µm.

### Movie S3

The motility of KIF1AMD-XKidSt was analyzed through time-lapse imaging recorded at 10 fps and played back at 25 fps. The dimensions of the displayed frame are 30 µm by 7 µm.

### Movie S4

The motility of KIF1A(1-393)LZ was analyzed through time-lapse imaging recorded at 10 fps and played back at 25 fps. The dimensions of the displayed frame are 30 µm by 7 µm.

### Movie S5

The motility of hKid-mNeonGreen (Green) and Cy3 labelled double-strand DNA (Red) was observed through time-lapse imaging recorded at 0.5 fps and played back at 10 fps. The dimensions of the displayed frame are 25 µm by 8 µm.

