## Supplemental Figures for "The chromokinesin Kid (KIF22) forms a homodimer, moves processively along microtubules and transports double-stranded DNA"

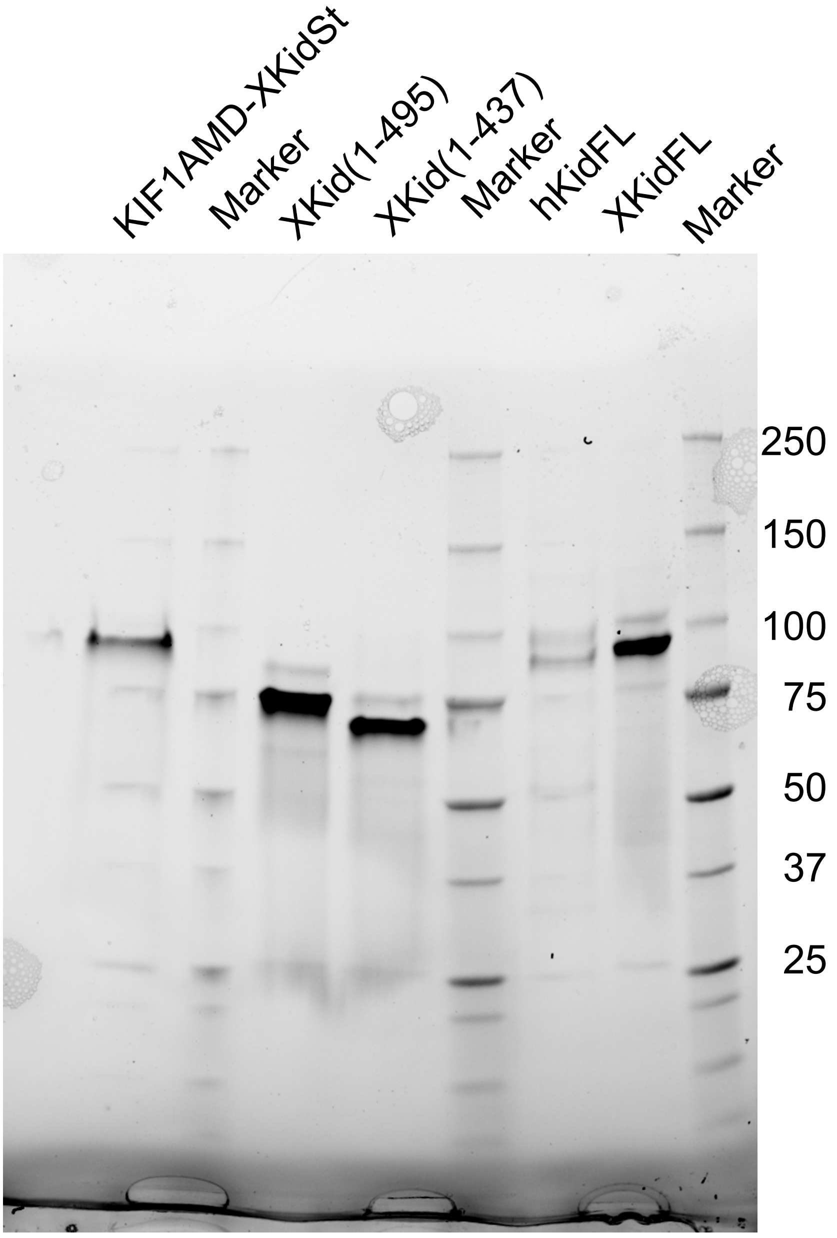


**Supplemental Figure S1 Full Scan Image of SDS-PAGE**

Purified proteins were separated by electrophoresis and visualized by Stain-Free gel (Bio-rad). The original image of Figures 1B, 3B and 4C. Numbers indicate molecular weight of the size marker.

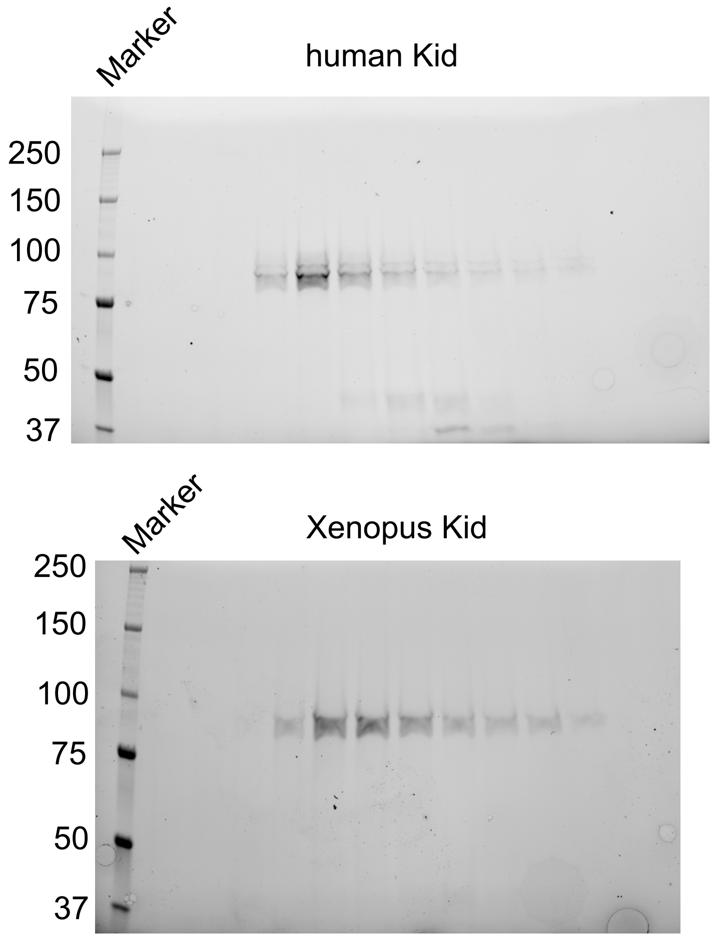


**Supplemental Figure S2 Full Scan Image of SDS-PAGE**

The original image of Figure 2A and B. Numbers indicate molecular weight of the size marker.

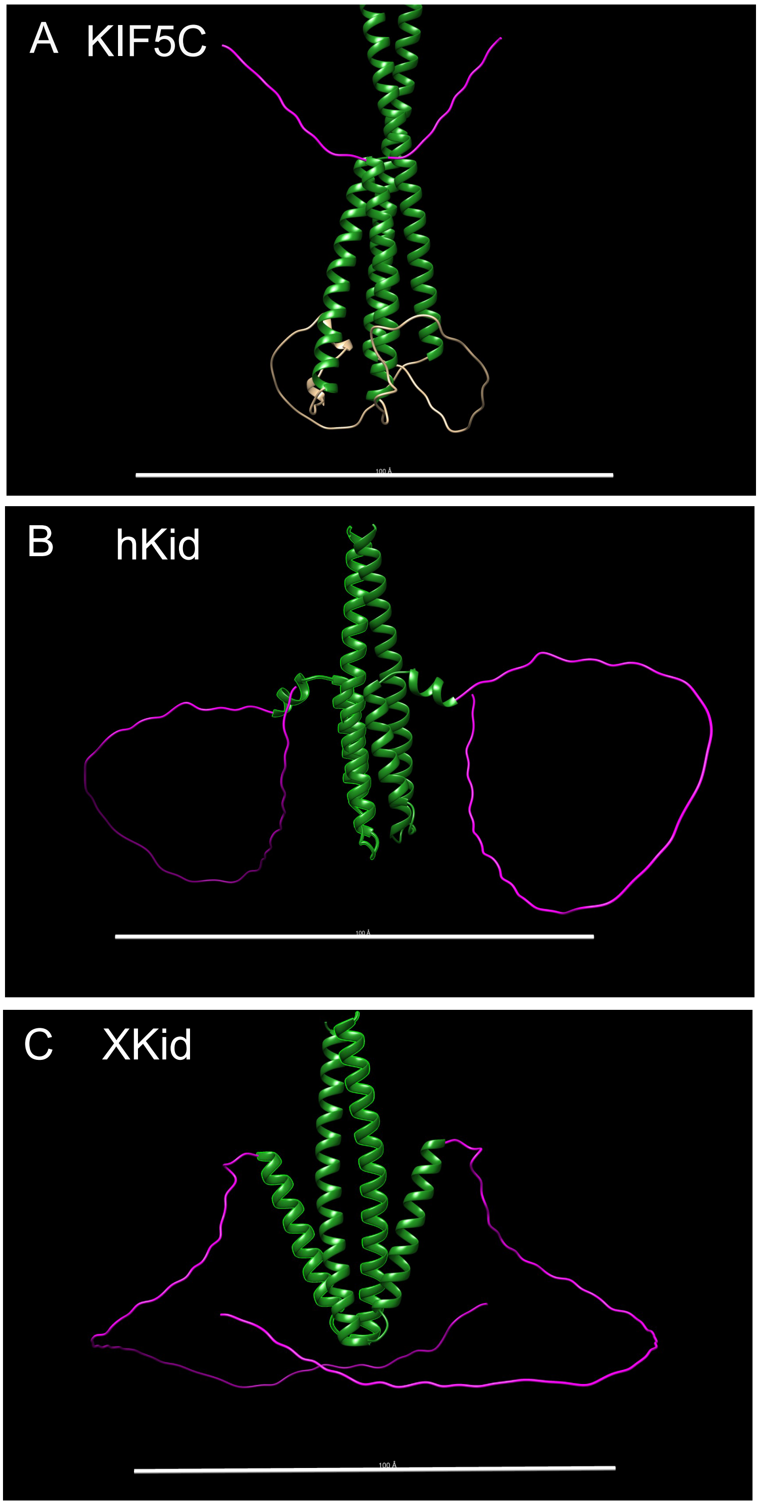


**Figure S3 Alphafold2 prediction of neck linker domain**

AlphaFold2 predictions were performed for sequences encompassing the region from the neck linker (highlighted in magenta) to the coiled-coil domain (colored in green). Relative to KIF5C (Panel A), both human Kid (hKid, Panel B) and Xenopus Kid (XKid, Panel C) exhibit extended flexible domains hypothesized to function as neck linkers. KIF5C is depicted in an autoinhibited conformation. The current study does not explore whether the folding structures in hKid and XKid represent an autoinhibited or active state, because the predictive analysis presented here is aimed at identifying potential neck linker and neck coiled-coil domains. Scale bars are set at 10 nm.

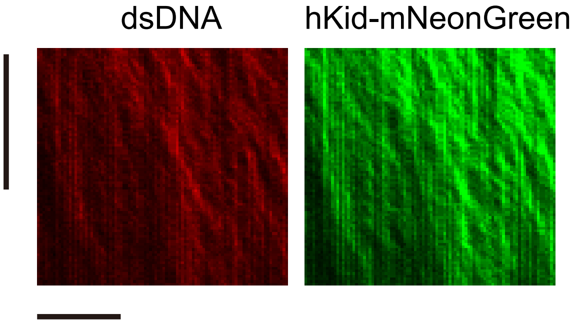


**Figure S4 hKid drive the movement of double-strand DNA**

Representative kymographs showing the movement of full-length hKid fused with mNeonGreen and double-strand DNA labelled with Cy3. Scale bars: horizontal, 5 μm; vertical, 5 minutes.
