## Supplementary material for "The chromokinesin Kid (KIF22) forms a homodimer, moves processively along microtubules and transports double-stranded DNA": Table S1

| name | sequence |
|---|---|
| hKidFL-mNG | atggccgcgggcggctcgacgcagcagaggcgacgcgagatggcggcagcttcagcggcggcgatctcaggagctggtcgctgtcggctaagcaagattggTgcAacGcgtcgtccacctccagctc<br>gcgtaagggtggctgtgcgactgcggccatttgtggatggaacagcgggagcaagtgatccccccctgtgtgcggggcatggacagctgctctctagagattgctaactggaggaaccaccaggagac<br>tctcaaataccagtttgatgccttctatggggagaggagtactcagcaggacatctatgcaggttcagtgcagcccatcctaaggcacttgctggaagggcagaatgccagtgtgcttgcctatgga<br>cccacaggagctgggaagacgcacacaatgctgggcagcccagagcaacctggggtgatcccgcgggctctcatggacctcctgcagctcacaagggaggagggtgccgagggccggccatgggccc<br>tttctgtcaccatgtcttacctagagatctaccaggagaaggtattagacctcctggaccctgcttcgggagacctggtaatccgagaagactgccggggggaatatcctgattccgggtctctccca<br>gaagcccatcagtagctttgctgattttgagcggcacttcctgccagccagtcgaaatcggactgtaggagccacccggctcaaccagcgctcctcccgcagtcatgctgtgctcctggtcaaggtg<br>gaccagcgggaacgtttggccccatttcgccagcgagagggaaaactctacctgattgacttggctgggtcagaggacaaccggcgcacaggcaacaagggccttcggctaaaagagagtggagcca<br>tcaacacctccctgtttgtcctgggcaaagtggtagatgcgctgaatcagggcctccctcgtgtaccttatcgggacagcaagctcactcgcctattgcaggactctctgggtggctcagcccacag<br>tatccttattgccaacattgcccctgagagacgcttctacctagacacagtctccgcactcaactttgctgccaggtccaaggaggtgatcaatcggccttttaccaatgagagcctgcagcctcat<br>gccttgggacctgttaagctgtctcagaaagaattgcttggtccaccagaggcaaagagagcccgaggccctgaggaagaggagatTgggagccctgagcccatggcagctccagcctctgcctccc<br>agaaactcagccccctacagaagctaagcagcatggacccggccatgctggagcgcctcctcagcttggaccgtctgcttgcctcccaggggagccaggggggcccctctgttgagtaccccaaagcg<br>agagcggatggtgctaatgaagacagtAgaagagaaggacctagagattgagaggcttaagacgaagcaaaaagaactggaggccaagatgttggcccagaaggctgaggaaaaggagaaccattgt<br>cccacaatgctccggcccctttcacatcgcacagtcacaggggcaaagcccctgaaaaaggctgtggtgatgcccctacagctaattcaggagcaggcagcatccccaaatgccgagatccacatcc<br>tgaagaataaaggccggaagagaaagctggagtccctggatgccctagagcctgaggagaaggctgaggactgctgggagctacagatcagcccggagctactggctcatgggcgccaaaaaatact<br>ggatctgctgaacgaaggctcagcccgagatctccgcagtcttcagcgcattggcccgaagaaggcccagctaatcgtgggctggcgggagctccacggccccttcagccaggtggaggacctggaa<br>cgcgtggagggcataacggggaaacagatggagtccttcctgaaggcaaacatcctgggtctcgccgccggccagcgctgtggcgcctcctGCGGCAGCGGTAGCgtgagcaagggcgaggaggata<br>acatggcctctctcccagcgacacatgagttacacatctttggctccatcaacggtgtggactttgacatggtgggtcagggcaccggcaatccaaatgatggttatgaggagttaaacctgaagtc<br>caccaagggtgacctccagttctcccccctggattctggtccctcatatcgggtatggcttccatcagtacctgccctaccctgacgggatgtcgcctttccaggccgccatggtagatggctccggc<br>taccaagtccatcgcacaatgcagtttgaagatggtgcctcccttactgttaactaccgctacacctacgagggaagccacatcaaaggagaggcccaggtgaaggggactggtttccctgctgacg<br>gtcctgtgatgaccaactcgctgaccgctgcggactggtgcaggtcgaagaagacttaccccaacgacaaaaccatcatcagtacctttaagtggagttacaccactggaaatggcaagcgctaccg<br>gagcactgcgcggaccacctacacctttgccaagccaatggcggctaactatctgaagaaccagccgatgtacgtgttccgtaagacggagctcaagcactccaagaccgagctcaacttcaaggag<br>tggcaaaaggcctttaccgatgtgatgggcatggacgagctgtacaagGGTTCTGGCTCTTGGAGTCATCCACAATTTGAGAAG |
| XKidFL-mSca | ATGGTTCTTACTGGGCCTCTCCAAAGAGAGTCGGTGAGTATGGCGAAGCGGGTGAGCATGTTGGATCAGCACAAGAAGTCCTCCTGTGCCCGGGTGCGTGTAGCCGTCAGACTGAGGCCCTACATGG<br>ATGAGAAGGATGAAGCTAAAGCGACCACAGTGTGTGTCAGGGGCCTGGACTCTCAGTCGCTGGAGATTGTCAACTGGAGAAACCAGCTGGAGACCATGCAGTACCAGTTTGATGCATTTTATGGGGA<br>CAGCGCCAGTCAGCGTGAGATCTACATGGGCTCTGTGTGCCACATTCTCCCGCACTTACTCATTGGCCAGAACGCCAGTGTTTTTGCTTATGGTCCCACGGGGGCAGGGAAAACTCACACAATGCTG<br>GGGAACCCCAATCAGCCTGGTGTGATCCCTCGGGCTGTTAGAGACTTGCTGCAGATGAGCCGGACAGCGGCCAGTGCCCCTGAGAATGAGAACTGGACTTATACCATAAACATGTCATATGTGGAGA<br>TTTACCAAGAGAAGGTCATGGATCTGCTGGAGCCAAAGAACAAAGACCTCCCCATCCGAGAGGACAAAGACCATAACATCCTGATCCCGGGCGTGACCCAAAAAATGATCAATTCTTTTGCAGATTT<br>CGATGAGCATTTTATCCCTGCCAGTCAGAACCGTACCGTGGCCTCTACGAAGCTCAATGACCGATCCAGTCGCAGCCACGCCGTGTTACTGATCAAGGTCCAGAAGAGTCAGCAGGTGGTACCATTT<br>CGGCAGCTGACTGGAAAGCTCTACCTGATAGACTTGGCTGGATCAGAAGATAATCGGCGCACTGGAAACCAAGGAATCCGACTAAAGGAGAGTGGAGCCATTAACTCCTCCTTATTCACGCTCAGCA<br>AAGTGGTGGATGCTCTGAACCAGGGGCTGCCCCGAATTCCATACAGAGACAGCAAGCTGACGAGACTGCTGCAGGATTCCCTGGGAGGAAGCGCCCACAGTGTTATGATCACTAACATCGCCCCAGA<br>GCAGACGTATTACTTCGACACATTGACGGCTCTGAACTTCGCTGCTAAATCCAAGCAGATCATTAACAAGCCTTTCAGCCAGGAAACCACCCAGACCGTGGTTCAGCCAGCCATGAAGAGGCCCAGG<br>GAGGAAACAGGCCACATAGCCGGTTCTCAGAAAAGAAAGAAATCCAAAAATGACTCCACTGAGTCGTCTCCCAATTCATCAATGGACACAGCGGGCAAACAGAAACTCAATTTGGCAACGCTGGATC<br>CCGCTGTGGTAGAGAGGCTGCTGAAACTGGATAAGATCCTGACCGAGAAGGGAAAGAAGAAGGCCCAGCTCTTGAGCACCCCCAAGAGAGAGCGAATGGCACTGCTGAAGAAATGGGAGGAAAGTCA<br>GATGGAGATCGAGAGACTAAAGGAGAAGCAAAAGGAGCTGGAGCAGAAAGCGATGGAAGCAGAGGCTCGACTGGAGAAATCCAATAACTCTGACCTGTCAGACTCCAGTGTCAGTGAAAACACTTTC<br>CGGGCCCCTCTCAGAGGCAGAAACACATCCACAGCGAAGGTCAAGAAAGTGCTGCGTGTACTGCCCATGCAGGGGAACAGCCAGTTACAAAGCACCGTAGAAGAGGGGATCCCCGTTTTTGAGAAGA<br>AGAAAAAGAAGAAACAGGTCACATGCGAGGGGCTTGAGAACCAGCCCACGTGGGAAATGAACATGAGGACAGACCTGCTTGAGAGCGGCAAGGAGAGAATCCTGAAACTACTCAACACGGGCTCAGT<br>AAAGGAACTGAAATCCCTGCAGAGGATCGGAGACAAGAAGGCCAAGCTGATTATTGGCTGGAGAGAAGTCAATGGGCCTTTTAAAAATGTGGAAGAGTTGGCGTGTTTGGAAGGAATCTCTGCTAAA<br>CAAGTATCGTCCTTTATAAAGGCAAATATCATGAGCAGCATCGCCAGCGGgcggccgctttcgaatATGGTCAGCAAAGGTGAAGCGGTAATAAAGGAGTTCATGCGCTTCAAGGTGCACATGGAAG<br>GTAGCATGAATGGACATGAGTTTGAGATTGAGGGAGAAGGTGAGGGCAGACCATACGAAGGAACACAGACGGCTAAGCTCAAAGTGACCAAGGGAGGCCCGTTGCCTTTCTCTTGGGACATTCTTAG<br>TCCCCAATTCATGTATGGAAGTAGAGCCTTCACGAAACATCCGGCGGATATACCCGATTATTACAAGCAGTCCTTTCCCGAGGGATTCAAATGGGAGCGTGTAATGAATTTTGAGGATGGAGGCGCA<br>GTAACAGTTACCCAAGACACTTCACTGGAGGATGGTACACTTATTTACAAAGTCAAACTTCGCGGCACTAATTTTCCGCCTGACGGTCCCGTTATGCAGAAAAAAACAATGGGATGGGAAGCCTCGA<br>CTGAACGTCTGTACCCTGAGGACGGCGTCCTGAAAGGTGACATTAAAATGGCTCTGCGCCTCAAAGACGGTGGACGTTACCTTGCTGACTTTAAAACTACTTACAAAGCAAAAAAACCGGTACAAAT<br>GCCCGGCGCGTACAACGTGGATCGCAAGTTGGACATTACTTCTCATAACGAGGATTACACCGTCGTTGAGCAGTATGAGAGGAGCGAAGGCCGCCATAGCACTGGAGGAATGGATGAATTGTATAAA<br>CTCGAGGGCAGTGGCAGTTGGTCGCATCCTCAATTCGAGAAGGGCGGCGGCAGTGGCGGCGGCAGTGGCGGCAGTGCCTGGAGTCATCCACAATTTGAGAAG |
| KIF1AMD-XKidSt | ATGGCCGGGGCTTCGGTGAAGGTGGCGGTGCGGGTCCGCCCCTTCAATTCCCGGGAAATGAGCCGTGACTCCAAGTGCATCATTCAGATGTCTGGAAGCACCACCACCATTGTTAACCCCAAACAGC<br>CCAAGGAGACGCCCAAAAGCTTCAGCTTTGACTACTCCTACTGGTCGCACACCTCACCTGAGGACATCAACTACGCGTCGCAGAAGCAGGTGTACCGGGACATCGGCGAGGAGATGCTGCAGCATGC<br>CTTTGAGGGATACAACGTGTGCATCTTCGCCTATGGGCAGACGGGTGCCGGCAAGTCCTACACCATGATGGGCAAGCAGGAGAAGGACCAGCAGGGCATCATCCCACAGCTCTGCGAGGACCTCTTC<br>TCTCGGATCAACGACACGACCAACGACAACATGTCCTACTCCGTGGAGGTCAGCTACATGGAGATTTACTGTGAGCGCGTCCGTGACCTCCTGAACCCCAAGAACAAGGGCAACCTTCGCGTGAGGG<br>AGCACCCACTGCTGGGGCCCTACGTGGAGGACCTCTCCAAGCTGGCTGTCACCTCCTACAATGACATCCAGGACCTCATGGACTCAGGGAACAAGGCCAGGACCGTGGCGGCCACCAACATGAATGA<br>GACCAGCAGTCGCTCCCACGCCGTCTTCAACATCATCTTCACCCAGAAGCGCCATGACGCAGAGACCAATATCACCACGGAGAAGGTGAGCAAAATCAGCCTGGTGGACCTGGCTGGGAGCGAGCGG<br>GCTGACTCCACGGGAGCCAAGGGCACGCGCCTCAAGGAGGGGGCCAACATCAACAAGTCGCTGACCACCCTGGGCAAGGTCATCTCCGCCCTGGCTGAAATGGACTCCGGACCCAACAAGAACAAGA<br>AAAAGAAGAAGACAGATTTCATTCCGTACCGAGATTCCGTGTTGACCTGGCTCCTCCGGGAAAACCTGGGCGGTAACTCAAGGACAGCTATGGTGGCAGCCTTGAGTCCTGCAGACATCAACTACGA<br>TGAGACCCTTAGCACGCTGAGGTATGCTGACCGGGCCAAGCAGATCATTAACAAGCCTTTCAGCCAGGAAACCACCCAGACCGTGGTTCAGCCAGCCATGAAGAGGCCCAGGGAGGAAACAGGCCAC<br>ATAGCCGGTTCTCAGAAAAGAAAGAAATCCAAAAATGACTCCACTGAGTCGTCTCCCAATTCATCAATGGACACAGCGGGCAAACAGAAACTCAATTTGGCAACGCTGGATCCCGCTGTGGTAGAGA<br>GGCTGCTGAAACTGGATAAGATCCTGACCGAGAAGGGAAAGAAGAAGGCCCAGCTCTTGAGCACCCCCAAGAGAGAGCGAATGGCACTGCTGAAGAAATGGGAGGAAAGTCAGATGGAGATCGAGAG<br>ACTAAAGGAGAAGCAAAAGGAGCTGGAGCAGAAAGCGATGGAAGCAGAGGCTCGACTGGAGAAATCCAATAACTCTGACCTGTCAGACTCCAGTGTCAGTGAAAACACTTTCCGGGCCCCTCTCAGA<br>GGCAGAAACACATCCACAGCGAAGGTCAAGAAAGTGCTGCGTGTACTGCCCATGCAGGGGAACAGCCAGTTACAAAGCACCGTAGAAGAGGGGATCCCCGTTTTTGAGAAGAAGAAAAAGAAGAAAC<br>AGGTCACATGCGAGGGGCTTGAGAACCAGCCCACGTGGGAAATGAACATGAGGACAGACCTGCTTGAGAGCGGCAAGGAGAGAATCCTGAAACTACTCAACACGGGCTCAGTAAAGGAACTGAAATC<br>CCTGCAGAGGATCGGAGACAAGAAGGCCAAGCTGATTATTGGCTGGAGAGAAGTCAATGGGCCTTTTAAAAATGTGGAAGAGTTGGCGTGTTTGGAAGGAATCTCTGCTAAACAAGTATCGTCCTTT<br>ATAAAGGCAAATATCATGAGCAGCATCGCCAGCGGgcggccgctttcgaatATGGTCAGCAAAGGTGAAGCGGTAATAAAGGAGTTCATGCGCTTCAAGGTGCACATGGAAGGTAGCATGAATGGAC<br>ATGAGTTTGAGATTGAGGGAGAAGGTGAGGGCAGACCATACGAAGGAACACAGACGGCTAAGCTCAAAGTGACCAAGGGAGGCCCGTTGCCTTTCTCTTGGGACATTCTTAGTCCCCAATTCATGTA<br>TGGAAGTAGAGCCTTCACGAAACATCCGGCGGATATACCCGATTATTACAAGCAGTCCTTTCCCGAGGGATTCAAATGGGAGCGTGTAATGAATTTTGAGGATGGAGGCGCAGTAACAGTTACCCAA<br>GACACTTCACTGGAGGATGGTACACTTATTTACAAAGTCAAACTTCGCGGCACTAATTTTCCGCCTGACGGTCCCGTTATGCAGAAAAAAACAATGGGATGGGAAGCCTCGACTGAACGTCTGTACC<br>CTGAGGACGGCGTCCTGAAAGGTGACATTAAAATGGCTCTGCGCCTCAAAGACGGTGGACGTTACCTTGCTGACTTTAAAACTACTTACAAAGCAAAAAAACCGGTACAAATGCCCGGCGCGTACAA<br>CGTGGATCGCAAGTTGGACATTACTTCTCATAACGAGGATTACACCGTCGTTGAGCAGTATGAGAGGAGCGAAGGCCGCCATAGCACTGGAGGAATGGATGAATTGTATAAACTCGAGGGCAGTGGC<br>AGTTGGTCGCATCCTCAATTCGAGAAGGGCGGCGGCAGTGGCGGCGGCAGTGGCGGCAGTGCCTGGAGTCATCCACAATTTGAGAAGTAA |
