## Supplementary material for "The chromokinesin Kid (KIF22) forms a homodimer, moves processively along microtubules and transports double-stranded DNA": Table S2

| protein | sequence |
|---|---|
| KIF5C_NL_NC_CC1 | M....(MD)...IKNTVSVNLELTAEEWKKKYEKEKEKNKTLKNVIQHLEMELNRWRNGEAVPEDEQISAKDQKNLEPCD<br>NTPIIDNIAPVVAGISTEEKEKYDEEISSLYRQLDDKDDEINQQSQLAEKLKQQMLDQDELLASTRRDYEKIQEELTRLQ<br>IENEAAKDEVKEVLQALEELAVNYDQKSQEVEDKTRANEQLTDELAQKTTTLTTTQRELSQLQELSNHQKKRATEILNLL<br>LKDLGEIGGI |
| XKid_NL_CC1 | M....(MD)...IINKPFSQETTQTVVQPAMKRPREETGHIAGSQKRKKSKNDSTESSPNSSMDTAGKQKLNLATLDPAV<br>VERLLKLDKILTEKGKKKAQLLSTPKRERMALLKKWEESQMEIERLKEKQKELEQKAMEAEARLEKSNN |
| hKid_NL_CC1 | M....(MD)...VINRPFTNESLQPHALGPVKLSQKELLGPPEAKRARGPEEEEIGSPEPMAAPASASQKLSPLQKLSSM<br>DPAMLERLLSLDRLLASQGSQGAPLLSTPKRERMVLMKTVEEKDLEIERLKTKQKELEAKMLAQKAEEKEN |
