## Supplementary material for "The chromokinesin Kid (KIF22) forms a homodimer, moves processively along microtubules and transports double-stranded DNA": Table S3

| protein | sequence |
|---|---|
| **DNA binding domain of XKid** | EGLENQPTWEMNMRTDLLESGKERILKLLNTGSVKELKSLQRIGDKKAKLIIGWREVNGPFKNVEELACLEGISAKQVSSFIKANI MSSIAS |
| **DNA1** | 5'-AGTCGATGCTACGTA-3' |
| **DNA2** | 5'-TACGTAGCATCGACT-3' |
